# Parenchymal precursor cells act in concert with canonical neural progenitors to facilitate neural circuit recovery following neuronal loss

**DOI:** 10.64898/2026.04.10.717664

**Authors:** William C. Tucker, Susanna L. Shepard, Paige E. Chambers, Anya Majji, John M. Boyd, Tracy A. Larson

## Abstract

Seasonal environmental variation represents one of the most pervasive and recurring forces shaping the vertebrate brain through cycles of neural circuit plasticity and remodeling that ultimately serves to maintain adaptive behavior. Although the contribution of adult neurogenesis to such remodeling is well-characterized across vertebrates, the role of astrocytes in supporting or enabling this process remains almost entirely unknown. Here, using the robust and natural seasonal loss of neurons within the sensorimotor nucleus HVC of the songbird, we identify a rapid astrocytic turnover event following neuronal loss and reveal a previously undescribed cellular mechanism supporting neural circuit recovery. Using lineage-specific and proliferation labeling, we establish the presence of a previously undescribed SOX2-positive neural progenitor-like population residing within the vertebrate parenchyma, outside the canonical ventricular zone niche. These parenchymal astrocyte precursor cells (pAPCs) maintain quantifiable, steady-state proliferation under homeostatic conditions, yet expand in pool size following seasonally-driven neuronal loss. The coordinated response of canonical neural progenitor cells and pAPCs produces new astrocytes that persist throughout the re-establishment of homeostasis. These findings reveal extensive astrocyte plasticity in the adult vertebrate brain, operating across both canonical and non-canonical progenitor niches, and challenge the prevailing view that parenchymal astrocytes are largely post-mitotic outside of injury contexts. More broadly, our findings establish a framework for understanding how glial populations contribute to neural circuit resilience both to the physiological and neural changes that recurring seasonal transitions drive, and to the accelerating environmental changes that increasingly challenge the adaptive capacity of animal brains worldwide.

## INTRODUCTION

Adult neurogenesis, or the generation of new neurons in the adult vertebrate brain, plays a vital role in learning, memory, and emotional regulation and allows the brain to respond to environmental changes, disease, and injury [1–4]. In vertebrates, adult-born neurons are generated by proliferative neural progenitor cells (NPCs) typically residing in specified niches along the ventricles of the brain. These NPCs, particularly radial glia, are of astrocytic-origin [5,6] and contribute not just new neurons, but also astrocytes and oligodendrocyte precursors in the adult brain [7–9]. The glia born in the adult brain via ‘gliogenesis’ regulate neurotransmitter and ion levels, provide metabolic support to neurons, contribute to the blood-brain barrier, and generally modulate and contextualize neural signaling [10,11]. Beyond homeostatic maintenance, astrocytes derived through adult gliogenesis also contribute to repair after neuronal injury and death [12,13]. Understanding the mechanisms regulating NPC proliferation, progeny fate specification, and astrocytic integration is therefore essential for understanding how neural circuits and the behaviors they support are maintained both under stable conditions and following environmentally-driven neural perturbation.

Seasonal environmental variation represents one of the most salient and recurring ecological forces shaping the vertebrate brain. Photoperiod, temperature, and hormonal cycles drive substantial remodeling of neural circuits across species, yet the cellular mechanisms enabling circuit recovery and behavioral maintenance across these transitions remain incompletely understood. Songbirds offer a uniquely tractable model for examining how brains respond, across levels of biological organization, to environmentally-driven neural change. In contrast to mammals, in which adult neurogenesis is restricted to two regions of the adult brain and occurs at relatively limited rates [14–16], songbirds exhibit quantifiably higher levels of adult neurogenesis across the entire telencephalon [17–20]. In Gambel’s white-crowned sparrow (*Zonotrichia leucophrys gambelii*) the sensorimotor region that controls singing behavior, called HVC (proper name [21]), functionally incorporates around 30,000 new neurons into an existing pool of around 90,000 neurons as sparrows enter each annual breeding season (Figure 1A and B) [22–24]. These new HVC neurons send projections over four millimeters away to the robust nucleus of arcopallium (RA; Figure 1A and B) [19, 25], and support the production of breeding song [26] for attracting mates and maintaining territories. As sparrows transition out of the breeding season, a sharp decline in circulating testosterone triggers the rapid death an equivalent number of HVC neurons, stimulating a reactive increase in NPC proliferation in the adjacent ventral ventricular zone (vVZ) and cessation of singing behavior [17,27–28]. This recurring, environmentally-coupled cycle of neuronal loss and regeneration makes the song control circuit an exceptional system for investigating how the tissues respond to seasonal perturbation.

**Figure 1.**
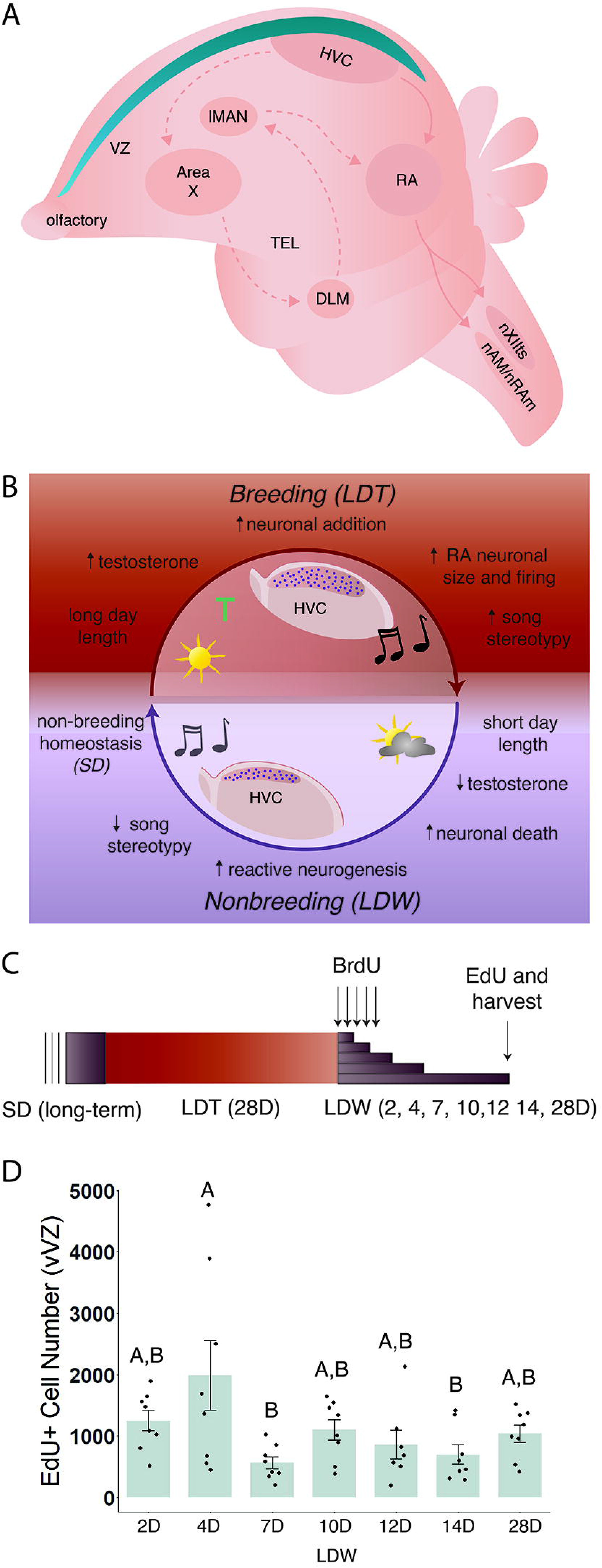
Gambel’s white-crowned sparrows exhibit extreme cyclical neuronal death that induces reactive proliferation of the nearby neural progenitor cells (NPCs) in the ventral ventricular zone (vVZ). (A) Simplified schematic of the avian brain with the descending motor pathway for singing behavior (solid lines) and the anterior forebrain pathway for learning of song (dotted lines). The classic NPC niche along the vVZ is highlighted in blue. Abbreviations: lMAN, lateral portion of the magnocellular nucleus of the anterior neostriatum; RA, robust nucleus of arcopallium; TEL, telencephalon; DLM, dorsolateral thalamic nucleus. (B) Cyclical regeneration and degeneration of HVC and the descending motor pathway with transition into and out of breeding condition, respectively. (C) Experimental design. (D) Reactive proliferation of NPCs in the vVZ. The number of EdU positive cells in the vVZ – representing NPC proliferation and their progeny generated in the previous two hours – increase significantly at two days following transition from breeding to nonbreeding condition. Data presented as mean ± SE, dots represent individual data with one dot per bird. Letters indicate significant differences across groups by post-hoc Tukey.

Despite extensive characterization of HVC neuronal dynamics [19,29–31], the astrocytes of HVC and the broader song control circuit have received remarkably little attention [32–34]. In the domesticated canary (*Serinus canaria domestica*), astrocyte numbers within HVC broadly mirror seasonal neuronal fluctuations across breeding and nonbreeding states [32]. Following stab injury to the zebra finch (*Taeniopygia guttata*) telencephalon, astrocytes upregulate androgen receptor expression [35], and following hippocampal injury, astrocytes express aromatase [36] and synthesize estrogens [37]. Together, these findings suggest that avian astrocytes participate in both homeostatic maintenance and injury-responsive repair similar to those of mammalian astrocytes, yet may exhibit a far greater degree of plasticity, consistent with the more extreme neurogenic responses observed seasonally in songbirds. The dramatic reactive NPC proliferation triggered by seasonal HVC neuronal death therefore offers a rare opportunity to examine astrocyte origin, turnover, and plasticity in direct relationship to an ecologically meaningful, environmentally-driven perturbation.

Here, we examine the fate of progeny born during seasonally-induced reactive proliferation of NPCs and identify a very rapid turnover event of astrocytes within HVC. We also characterize a previously undescribed population of SOX2-positive cells within the telencephalon that not only proliferate at steady levels during homeostasis, but also increase in number following extreme seasonal neuron death in HVC. Investigating the relative contribution of this novel population of parenchymal astrocyte precursor cells (pAPCs) compared to NPCs in the classic ventricular zone niche, we find that an increase in pAPCs corresponds to an increase in the number of newly-born astrocytes and neurons within HVC. Together our results suggest that reactive proliferation following environmentally-driven neuronal death occurs not only within the canonical NPC niche at the ventricle, but also more generally within the parenchyma of the avian brain. This coordinated proliferative response drives expansion of the pAPC pool, astrocyte turnover, and contributes to restoration of homeostasis following seasonal neural degeneration. Our findings reveal extensive, environmentally-coupled astrocyte plasticity and a previously undescribed proliferative cell population in the avian telencephalon, laying the foundation for understanding how glial cells and their precursors – both within and beyond canonical niches – enable neural circuit resilience in the face of recurring environmental and physiological change.

## RESULTS

### Confirmation of degeneration and reactive proliferation

To test the fate specification and survival of the progeny derived from seasonally-induced reactive NPC proliferation in the ventral ventricular zone (vVZ), we housed Gambel’s white-crowned sparrows in breeding-like conditions and then transitioned sparrows into non-breeding conditions to induce degeneration of HVC and its behavioral output, song, as previously described (Figure 1C)[17]. HVC morphology and neuron number were consistent with those of degenerated HVC as previously reported [17] with no significant differences across time points of degeneration (ANOVA, F(_6,57_)=1.6894; p=0.1402), F(_6,48_)=1.0290; p=0.4182, respectively; SI Table 1). Reactive proliferation was confirmed as an increase in EdU positive cells in the vVZ at two days post transition into non-breeding conditions with a return to basal levels by seven days, as previously reported [17] (Figure 1D; ANOVA, F(_6,48_)=3.1368; p= 0.0113, SI Table 1). Song was also observed in the expected pattern (Figure 2), with LDT birds singing more complete songs (ANOVA, F(_7,110_)=18.4631; p<0.0001, Figure 2C and SI Table 1). All together this data affirms that our experimental setup replicated the previously reported neural and behavioral plasticity following transition into nonbreeding conditions, allowing for deeper examination of reactive proliferation and progeny fate.

**Figure 2.**
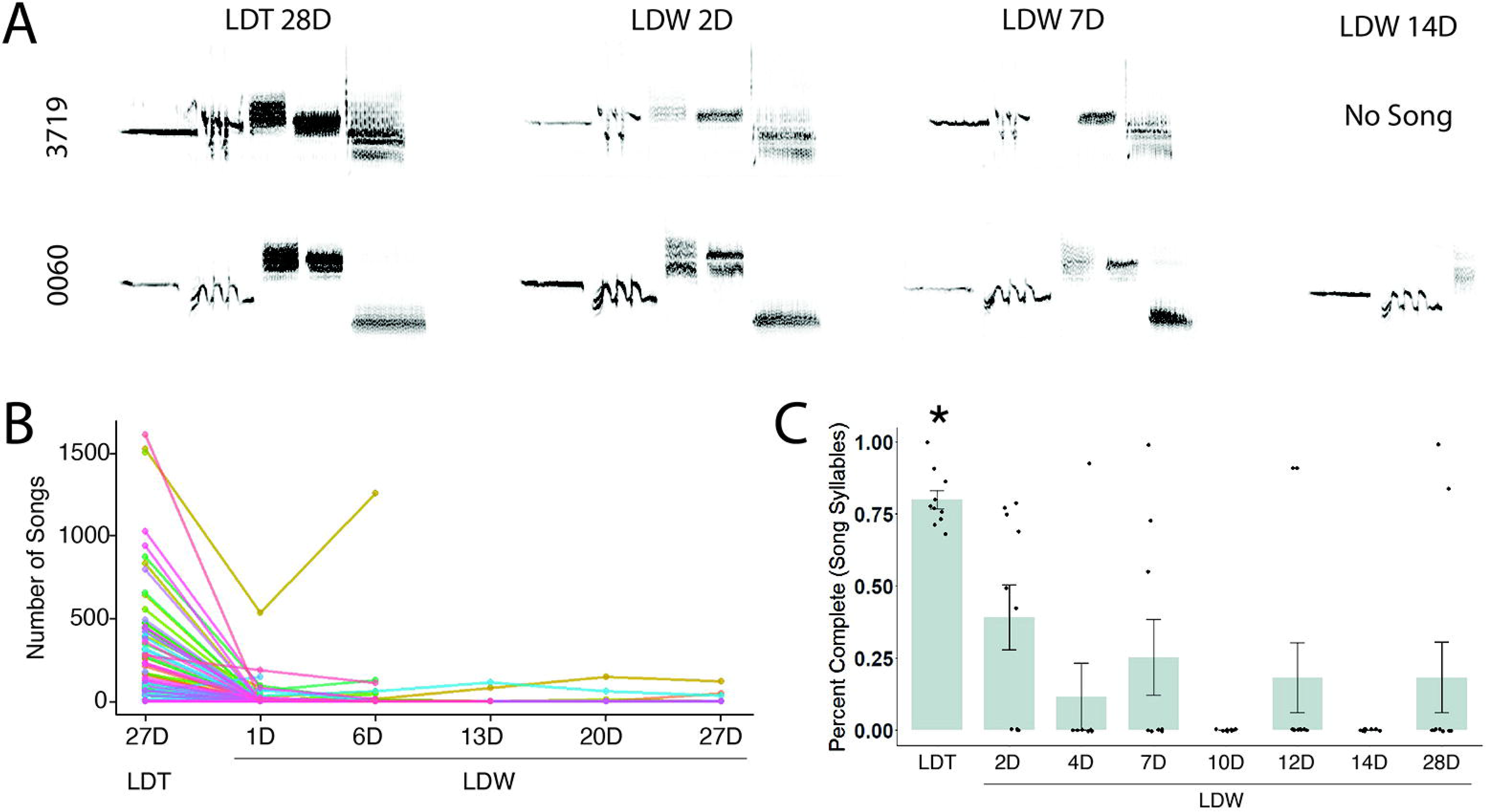
Singing quality and quantity decreases with transition out of breeding conditions. (A) Representative sonograms of two individual birds, 3719 and G0060, as each progressed through HVC degeneration and reactive proliferation. Song degrades in structure and completeness with time into non-breeding condition. (B) The number of songs sung per day by each experimental bird (n=74). Song rate rapidly decreases as soon as two days into non-breeding conditions. (C) Percent complete across experimental timepoints (n=9 to 11 per group). Birds in breeding conditions sang significantly more complete songs than birds in nonbreeding conditions, p<0.05 denoted with an asterisk (post-hoc Tukey).

### Birth and survival of neurons and astrocytes from reactive proliferation event

To identify the fate of progeny generated during seasonally-induced reactive proliferation, we labeled the progeny of proliferative cells using BrdU administered for five days during the long day withdraw (LDW)-associated reactive proliferative event (Figure 1C). Birds were subsequently housed in non-breeding LDW conditions for periods ranging from two to twenty-eight days, allowing recently born progeny to migrate and incorporate into HVC [38].

Quantification of the number of BrdU positive cells within HVC that co-labeled with the neuronal lineage-marker HuC/D (ELAV3/4; Figure 3A) revealed a significant increase in new-born HVC neuron number two days into HVC degeneration (Figure 3B and SI Table 2; ANOVA, F(_6,48_)=19.6345; p<0.0001). The number of BrdU and HuC/D positive cells decreased by four days and remained basal throughout the remainder of LDW. Likewise, quantification of new immature astrocytes labeled with vimentin (Figure 3A) identified a peak in newly generated astrocytes at two days into LDW that decreased to basal levels by four days (Figure 3C and SI Table 2; ANOVA, F(_6,46_)=12.3438; p<0.0001). The number of BrdU-labeled progeny of neuronal fate was consistently lower, between 20% and 34%, than those of astrocytic fate during both the peak and basal levels of NPC proliferation (Figure 3D and SI Table 2; ANOVA, F(_6,37_)=1.1241; p=0.3675). Together these results demonstrate that progeny of reactive NPC proliferation fate-specify into both neurons and astrocytes, with the majority of newly-born cells surviving long-term as astrocytes.

**Figure 3.**
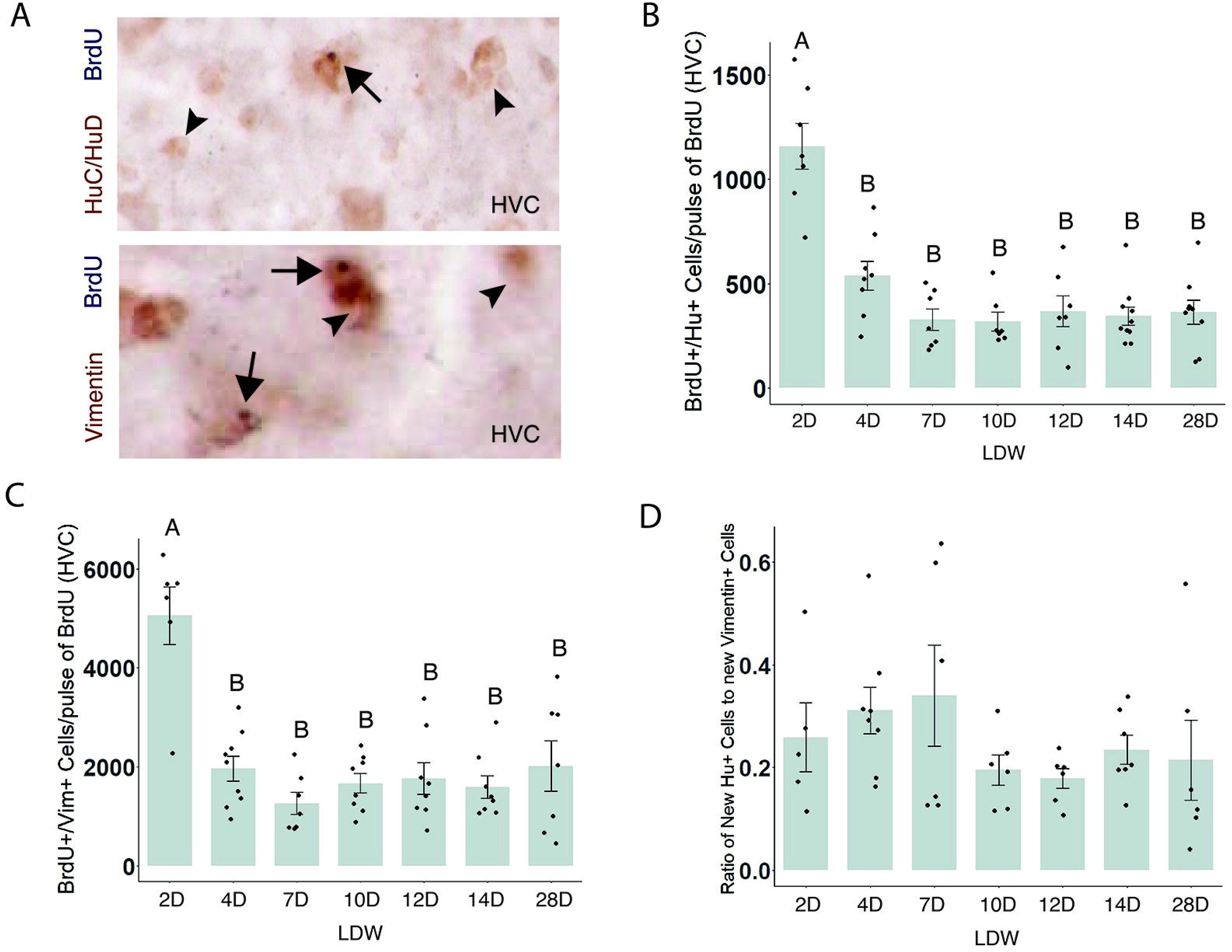
The specification and survival of cells born from reactive proliferation. (A) Representative image of new and pre-existing neurons and astrocytes in HVC. Neurons are stained with DAB in brown using a HuC/D antibody (top), astrocytes are stained with DAB in brown using a vimentin antibody (bottom). BrdU positive cells have DNA labeled by a BrdU antibody in black. Long arrow indicates BrdU positive cells. Short arrow indicates examples of pre-existing (non-BrdU positive) neurons/astrocytes within HVC. (B) The number of BrdU and HuC/D positive neurons born during reactive proliferation (n=7 to 11 per group) (C) The number of BrdU and vimentin positive astrocytes born during reactive proliferation (n=7 to 8 per group). (D) The ratio of new neurons to new vimentin positive astrocytes in HVC across song circuit regression (n=5 to 8 per group). (E) The proportion of BrdU-positive neurons to BrdU-positive astrocytes. All data presented as mean ± SE, dots represent individual data with one dot per bird. Letters indicate significant differences across groups by post-hoc Tukey.

### Low turnover of neurons, high turnover of astrocytes following HVC degeneration

To further assess the proportional survival of new cells from each fate generated as a result of reactive proliferation, we calculated the proportion of BrdU-labeled neurons and astrocytes relative to their respective peaks at two days. For neurons, 46.4% percent of BrdU-labeled HuC/D neurons survive to four days with 31.2% surviving to twenty-eight days (Figure 4A and SI Table 2; ANOVA, F(_5,39_)=1.886; p=0.1190). Neurons generated during the reactive proliferation event comprised 3.95% at the peak and 0.93% to 1.57% of the total population throughout the remainder of LDW (Figure 4B and SI Table 2; ANOVA, F(_6,46_)=13.8425; p<0.001). The significant decrease in number of BrdU-labeled neurons at four days and the decrease in proportion of neurons from the total neuronal population by ten days suggests that: 1) the survival of new neurons declines rapidly and continues to decline throughout HVC degeneration and return to homeostasis, and 2) the birth of new neurons from reactive proliferation do not contribute to a substantial neuronal turnover event.

**Figure 4.**
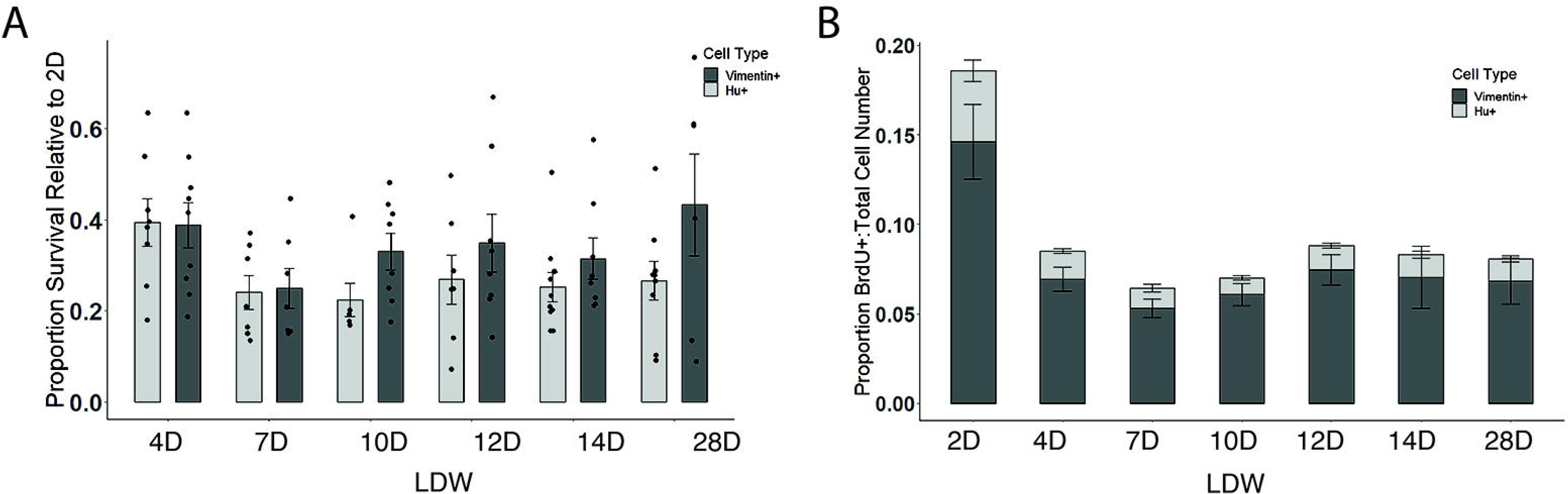
Survival and turnover of neurons and astrocytes generated during reactive proliferation. (A)The proportion of new neurons (light grey) and astrocytes (dark grey) relative to their respective total pool within HVC. The proportion of new vimentin positive astrocytes to the total population is higher than the proportion of new neurons added to HVC. (B) The proportion of new BrdU-positive cells co-labeled with HuC/HuD in light grey stacked on the proportion of BrdU-positive cells co-labeled with vimentin in dark grey. Data presented as the number of co-labeled BrdU cells at four or 28 days divided by the number co-labeled at two days. New neurons and astrocytes are rapidly culled with a proportion surviving for at least 28 days following birth.

Assessing astrocyte survival, we found that 38.8% percent of BrdU-labeled vimentin-positive astrocytes persist to four days and 43.2% persist to twenty-eight days (Figure 4A and SI Table 2; ANOVA, F(_5,40_)=1.0258; p=0.4156). Of the total astrocyte population, astrocytes generated during reactive proliferation represented 14.60% at the peak and between 5.30% to 7.47% throughout the remainder of LDW (Figure 4B and SI Table 2; ANOVA, F(_6,46_)=6.1529; p<0.0001). The lack of a significant decrease in survival of BrdU-labeled astrocytes across the time course of degeneration in combination relative to two days with the relatively higher (than neuron) proportion of new immature astrocytes in the total astrocyte population, suggests that a large number of astrocytes born during reactive proliferation incorporate into HVC and represent a substantial astrocytic turnover event.

### Identification of a population of proliferative SOX2-positive cells in the parenchyma

We aimed to determine whether the decrease in the number of newly-born vimentin positive astrocytes (Figure 3C) was due to culling or maturation (and therefore down regulation of vimentin protein). To do so, we immuno-labeled for BrdU with the mature astrocytic maker, glial fibrillary acidic protein (GFAP) across the same time course. To our surprise, rather than finding a gradual increase in GFAP positive cells co-labeled with BrdU as vimentin and BrdU positive cell numbers declined, we found a significant uptick in GFAP cells co-labeled with BrdU at two days (Figures 5A and B, SI Table 3; ANOVA, F(_6,47_)=12.0969; p<0.0001). This increase in newly-born GFAP positive cells at two days occurs at a timepoint at which incoming vVZ-derived cells would likely be unable to migrate from the vVZ, nor mature following birth, which is reported to take 1-2 weeks at minimum [39]. This raised the possibility that perhaps these GFAP-positive astrocytes were proliferative cells themselves or products of local proliferation outside of the VZ rather than mature quiescent astrocytes. To test the possibility that progenitor-like cells might exist and proliferate within the avian parenchyma and specifically within HVC, we labeled tissue from the same experimental birds with SOX2, a routinely-used lineage marker of proliferative neural progenitor cells [40]. After confirming SOX2 labeling of NPCs in the classic VZ niche (Figures 5C and D), we found that, indeed, there were cells positive for SOX2 within HVC (Figures 5C and E) and the telencephalon more broadly (Figures 5C and F).

**Figure 5.**
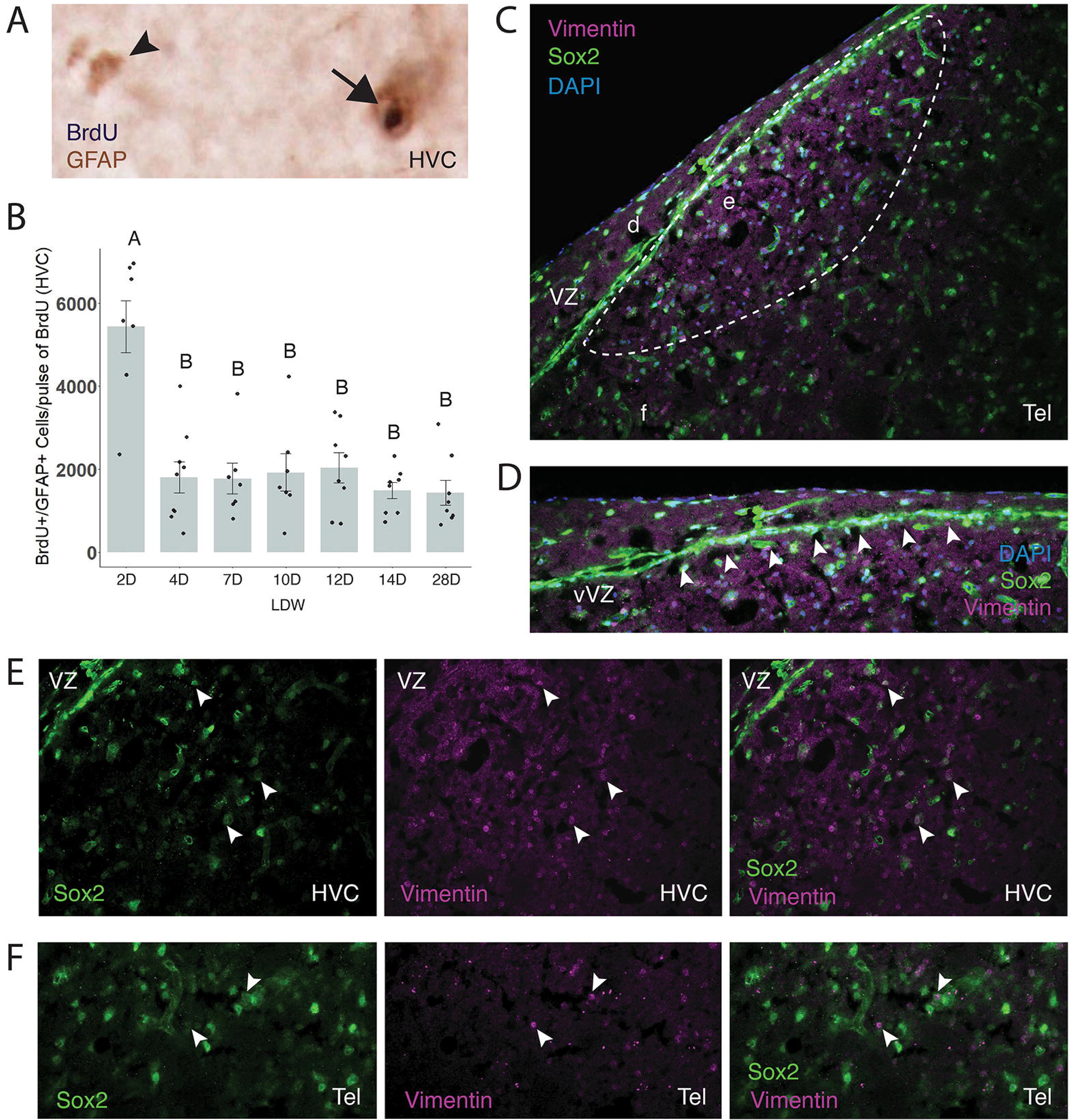
A previously undescribed population of proliferative progenitor-like cells within the parenchyma of the avian brain. (A) Tissue labeled for BrdU and GFAP, showing new GFAP positive cells within HVC. Black arrow indicates a GFAP positive cell with BrdU positive nucleus, arrowhead indicates a GFAP positive cell with no BrdU label. (B) The number of GFAP positive astrocytes within HVC co-labeled for BrdU across the time course of HVC degeneration and reactive proliferation. The number of GFAP cells co-labeled with BrdU in HVC rapidly peaks at two days, suggesting in-place proliferation of GFAP positive cells. Data presented as mean ± SE, dots represent individual data with one dot per bird. Letters indicate significant differences across groups by post-hoc Tukey. (C) Representative z-stack image (20x) of avian telencephalon labeled with fluorescent anti-SOX2 and anti-vimentin revealing a substantial population of SOX2-positive cells within the vVZ, HVC and the surrounding telencephalon. Lower case letters indicate location of zoomed-in images in D-F. (D) Z-stack image (60x) of the vVZ just above HVC showing the expected pattern of SOX2 labeling in the classic stem and progenitor cell niche at the vVZ. (E) Z-stack image (60x) of HVC revealing SOX2-positive cells (green) co-labeled with vimentin (purple; arrowheads) within HVC specifically. (F) Z-stack image (60x) of the surrounding telencephalon revealing SOX2-positive cells co-labeled with vimentin (arrowheads) within the parenchyma generally.

The presence of SOX2-positive cells beyond the VZ suggested that perhaps a previously undescribed population of progenitor-like cells in the parenchyma existed in the avian brain and maybe even contributed to reactive proliferation. Thus, to investigate whether or not these SOX2-positive cells within the parenchyma were progenitor cells, we assessed the following criteria for any progenitor cell: 1) proliferative potential, 2) capability of self-renewal, and 3) capacity for generating progeny that differentiate into non-self cells. First, to confirm that these SOX2-positive cells in the parenchyma are capable of proliferation, we co-labeled SOX2-positive cells with EdU that had been administered two hours prior to tissue harvesting for labeling active cellular proliferation. We identified SOX2-positive cells across the telencephalon that co-labeled with EdU (Figure 6A) including HVC and a target nucleus of HVC called the robust nucleus of arcopallium (RA), which has been reported to not incorporate adult-born neurons (Figure 6B) [19]. Quantification of SOX2-positive cells that recently proliferated and incorporated the thymidine analog EdU within the past two hours of the experiment revealed a substantial population of proliferating SOX2-positive cells within HVC, the rate of which was stable across the experimental time course (Figure 6C and SI Table 3, ANOVA, F(_6,45_)= 0.3355; p=0.9146).

**Figure 6.**
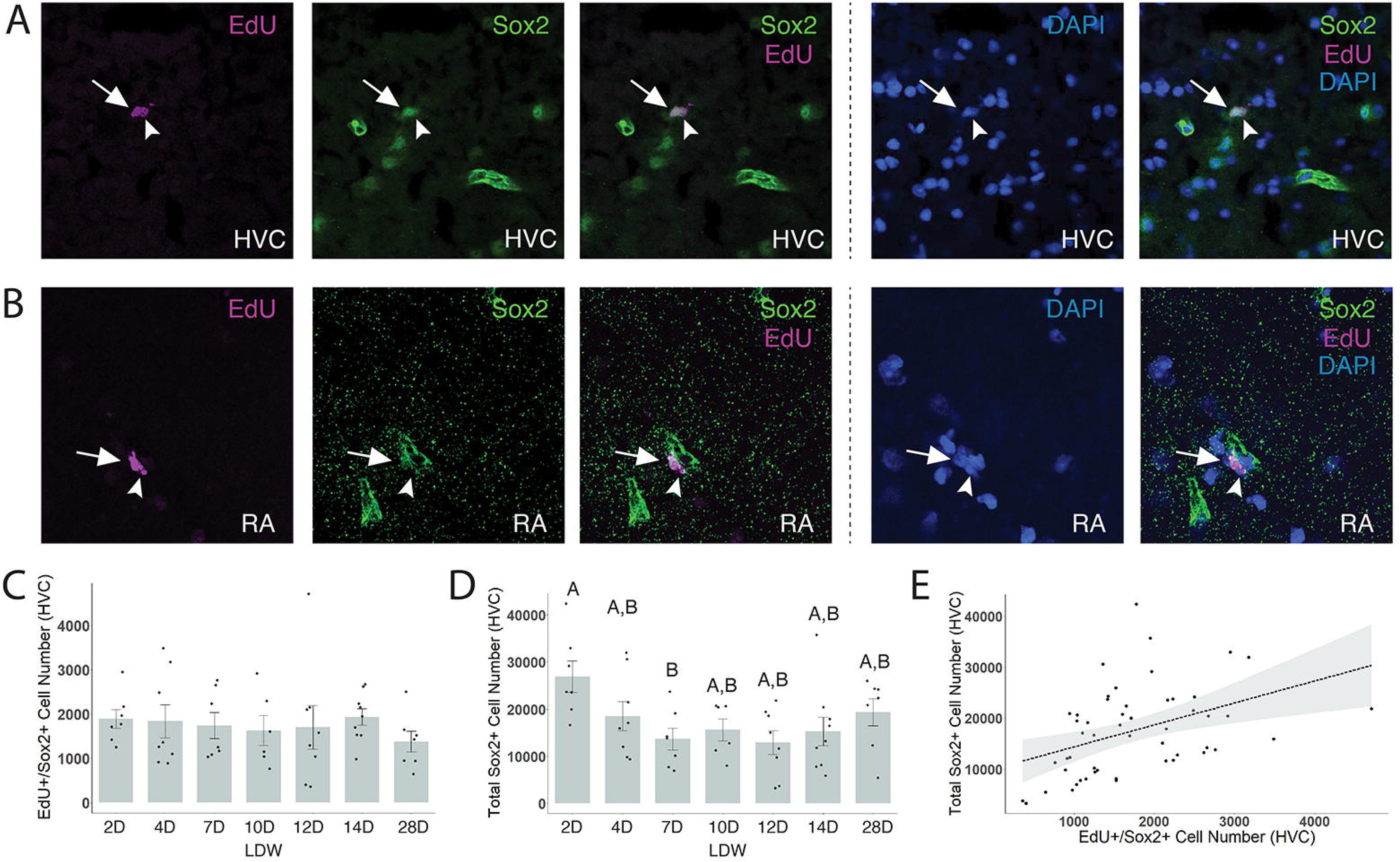
SOX2-positive cells located outside the classic VZ niche are capable of proliferating. Representative z-stack image (60x) of tissue containing (A) HVC and (B) RA labeled with anti-SOX2 immunofluorescence (green) and EdU Click-It chemistry (purple). SOX2-positive cells within HVC and RA label with EdU that had been administered two hours prior to tissue collection. (C) The number of EdU and SOX2 co-labeled cells within HVC exist at a sizable, non-zero population that remains stable throughout HVC degeneration. (D) The total number of SOX2-positive cells within HVC are significantly higher at two days following transition into nonbreeding conditions that other time points into HVC degeneration and return to homeostasis. The larger pool of the SOX2-positive cells within HVC suggests that these proliferative cells self-renew and are responsive to local conditions. (C-D) Data presented as mean ± SE, dots represent individual data with one dot per bird. Letters indicate significant differences across groups by post-hoc Tukey. (E) Pearson’s correlation between total SOX2-positive cells and SOX2-positive cells co-labeled with EdU within HVC (n=46; r^2^=0.2325, p<0.0001).

### Self-renewal of parenchymal neural progenitor cells

We next sought to determine whether these SOX2-positive cells were capable of self-renewal by quantifying the total population size of SOX2-positive cells across the time course of HVC degeneration and return to homeostasis. We found that the total number of SOX2-positive cells within HVC increased significantly at two days following transition into nonbreeding conditions and returned to baseline by seven days (Figure 6D and SI Table 3; ANOVA, F(_6,45_)= 2.7007; p=0.0252). We found a significant positive correlation between the total number of SOX2-positive cells and those co-labeled with EdU (Figure 6E and SI Table 4; r^2^=0.2325, p<0.0001), further suggesting that this population is self-renewing.

To further clarify whether the uptick in GFAP-positive cells also positive for BrdU at two days into HVC degeneration represented local expansion of the putative parenchymal astrocyte precursor pool rather than rapid maturation of astrocytic progeny (aside from physical location), we performed a Pearson’s correlation between GFAP and BrdU co-labeled cells and the total number of SOX2-positive cells within HVC. We found a significant positive correlation between the two (SI Table 4; r^2^=0.2798, p<0.0001), supporting the hypothesis that the GFAP positive cells co-labeled with BrdU represent a significant portion of the proliferative SOX2-positive cell pool within HVC.

### Progeny of parenchymal neural progenitor cells

Finally, to assess whether or not these SOX2-positive cells could have the potential to generate adult-born neurons and astrocytes, we identified relationships between SOX2-positive cells and fate-specified cell populations using Pearson’s correlations. We found a significant positive correlation between new BrdU/vimentin and EdU/SOX2-positive cells (Figure 7A and SI Table 4; r^2^=0.0923, p=0.0358). We also found a significant correlation between the number of new BrdU/vimentin and total number of SOX2-positive cells (Figure 7B and SI Table 4; r^2^=0.3282, p<0.0001). Analyzing the relationship between SOX2-positive cells and new HuC/D neurons in HVC, we found a significant positive correlation (Figure 7C and D, SI Table 4; r^2^=0.1719 p=0.0020), but no significant correlation between new neurons and the SOX2 cells co-labelled with EdU (Figure 7B and C, SI Table 4; r^2^=0.0002, p=0.9270).

**Figure 7.**
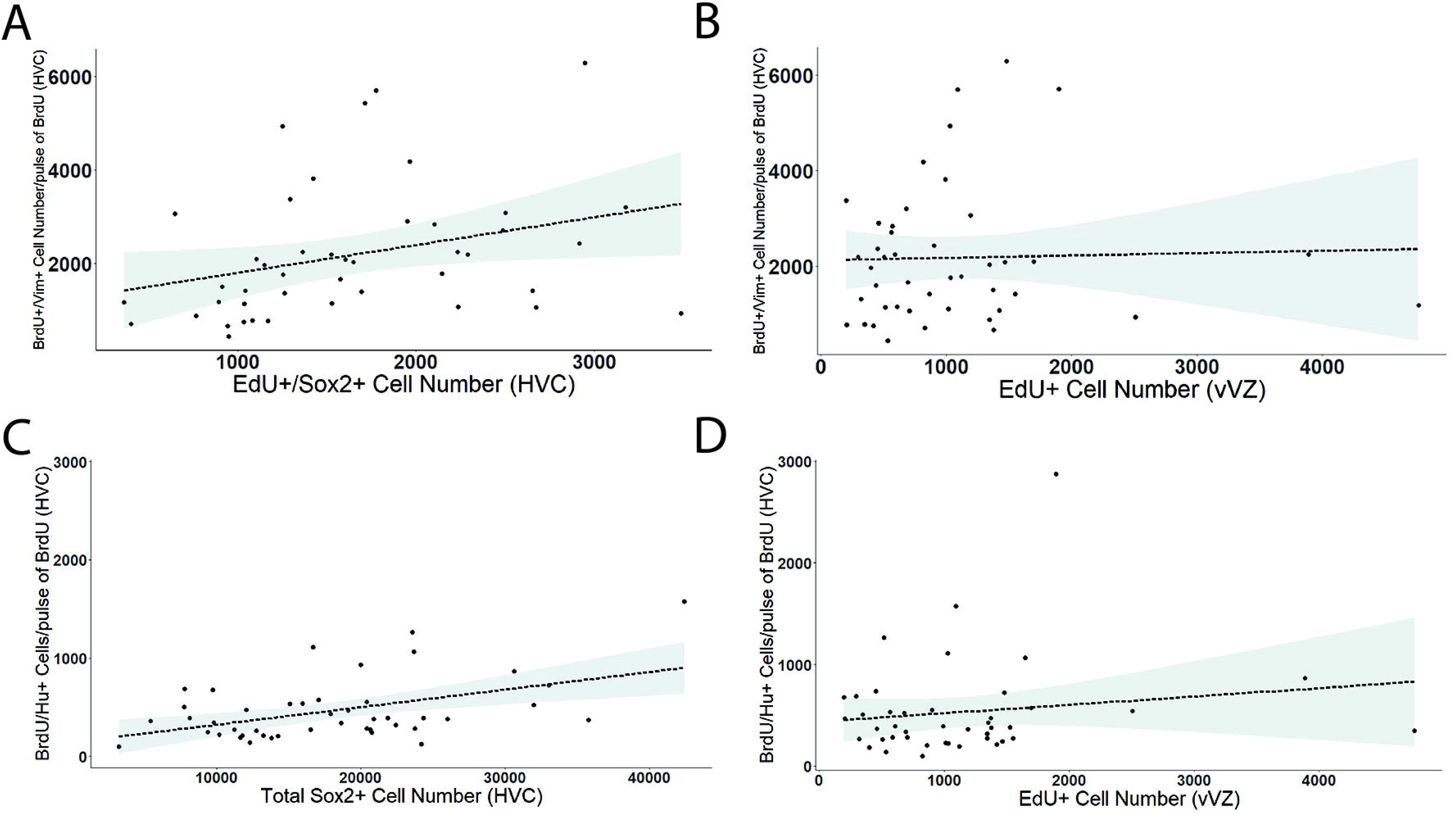
Total number and proliferation of SOX2-positive cells in the parenchyma but not EdU positive cells in the vVZ correlate with recently-born vimentin positive and HuC/D positive cells. (A) Pearson’s correlation of EdU positive and SOX2-positive cell number to BrdU positive and vimentin positive cell number (n=43; r^2^=0.2325, p<0.0001). (B) Pearson’s correlation of EdU positive cells in the vVZ to BrdU/vimentin positive cell number (n=43; r^2^=0.0012, p=0.8150). (C) Pearson’s correlation of BrdU/Hu positive cells to the total SOX2 population (n=44; r^2^=0.1712, p=0.0020). (D) Pearson’s correlation of BrdU/Hu positive cells to EdU positive number in the vVZ (n=42; r^2^=0.0485, p=0.1243).

To identify the relative contribution of NPC proliferation in the vVZ to astrocyte and neuronal addition in HVC, we examined the relationship between EdU positive cell number in the vVZ (the canonical niche) and total vimentin positive cell number in HVC and found no significant relationship (Figure 7B and SI Table 4; r^2^=0.0003, p=0.6935). Likewise, we found no significant correlation between EdU positive cell number in the vVZ and HuC/D positive neurons in HVC (Figure 7D and SI Table 4; r^2^=0.0485, p=0.1243). Finally, we examined the relationship between vVZ proliferation and the SOX2 pool and found no relationship between the number of EdU labeled cells in the vVZ and the total SOX2 population (SI Table 4; r^2^=0.0037, p=0.6775) nor SOX2-positive cells co-labeled with EdU (SI Table 4; r^2^=0.0187, p=0.3438).

Together these results suggest that rapid proliferation of progenitor cells in HVC, but not the more-classic progenitors in the VZ proliferation, drive a significant proportion of astrocyte and neuronal addition in HVC immediately following degeneration. All together our data suggest that these newly-identified Sox-2 positive cells are: 1) proliferative, 2) capable of self-renewal, and 3) contributing new fate-specified astrocytes and neurons within HVC. Satisfying these criteria of defining a progenitor cell establishes these SOX-2 positive cells as a newly identified population of precursor cells in the parenchyma of the avian brain – here termed parenchymal astrocyte precursor cells (pAPCs) – that are responsive to natural, seasonally-induced degeneration of local neuronal populations.

## DISCUSSION

We examined the fate of progeny born during seasonally-induced proliferation of vVZ-residing NPCs in the songbird telencephalon and identified a rapid astrocyte turnover event within HVC following environmentally-driven neuronal loss. Establishing the existence of a previously undescribed SOX2-positive NPC-like population within the parenchyma of the telencephalon, we found that these parenchymal astrocyte precursor cells (pAPCs) proliferate at steady rates during homeostasis and significantly expand their pool size following extreme seasonal neuronal death in HVC. Together these results demonstrate that reactive proliferation of canonical vVZ NPCs and pAPCs drives the rapid expansion and then turnover of the HVC astrocyte population, enabling return to homeostasis and perhaps even circuit regrowth the subsequent breeding season.

### Turnover of astrocytes and neurons following environmentally-driven neural degeneration

Adult songbirds exhibit robust and ongoing neuronal turnover [17–19], with new neurons arising from NPCs in the canonical ventricular niche [41–42] in a manner that is seasonally modulated in many species of songbirds [22,43–44]. Seasonal neurogenesis is driven by environmental cues, primarily photoperiod and the downstream hormonal cascades it triggers, making songbird HVC a powerful natural model for studying how brains respond to recurring ecological change across levels of biological organization. Although astrocyte number had been previously shown to fluctuate seasonally [32] and following neural injury (discussed below) in the songbird brain, the baseline rate of astrocyte turnover and the time course of seasonally-induced turnover had not been quantified prior to this study.

Inducing seasonal degeneration of HVC in male white-crowned sparrows rapidly increased the number of newly-born vimentin-positive astrocytes (Figure 3C). The number of new vimentin-positive cells subsequently declined by seven days to approximately 25% of peak values, recovering to around 43% of peak values by twenty-eight days (Figure 4A and SI Table 2). This apparent increase in proportional survival at twenty-eight days likely reflects two concurrent processes: first, the maturation of astrocytes born from reactive proliferation in the canonical vVZ niche [17]; and second, the accumulation of immature astrocytes born from both reactive and homeostatic proliferation of the newly-described pAPCs at two days and beyond (discussed further below). Regardless of origin, newly-born astrocytes persisting after HVC degeneration represented between five and seven percent of the total astrocytic population at twenty-eight days (Figure 4B and SI Table 2). The overall decline in BrdU-labeled vimentin-positive astrocytes across the time course suggests that remaining newly-born astrocytes either failed to integrate into HVC and died, transitioned to a mature state in which vimentin expression is downregulated [32,45], or some combination of both. To precisely discriminate between these possibilities, quantification of astrocytes bearing mature markers such as GFAP would, in principle, allow estimation of survival rates following reactive proliferation. Pursuing this approach, we found that 5.5% of GFAP-positive astrocytes within HVC were BrdU-labeled at twenty-eight days (SI Table 2); however, this result should be interpreted cautiously as BrdU- positive GFAP-expressing cells could represent either mature surviving astrocytes or slowly-dividing parenchymal precursor cells that retained BrdU label. Furthermore, given the dynamic nature of this cell population, there is likely some degree of marker overlap between vimentin and GFAP during astrocyte maturation [45] that was not quantified here. Resolving the precise rates of astrocyte survival versus BrdU retention in slow-cycling pAPCs, and clarifying the functional overlap between vimentin- and GFAP-expressing astrocyte populations across maturation states, will require dedicated fate-mapping analyses in future studies.

In contrast, neurons born from seasonally-induced reactive proliferation failed to survive long-term, as anticipated given HVC neuron numbers remain low throughout the non-breeding season. Approximately 25% of neurons born and surviving to two days following onset of HVC degeneration were still present at twenty-eight days, the time point for return to non-breeding homeostasis (Figure 4A and SI Table 2). Of the total population of neurons, this newly-born and surviving population represented only around 1% of the total neuronal population (Figure 4B and SI Table 2). These results are broadly consistent with previous studies reporting low baseline daily neuronal addition rates in other songbird species: approximately 0.5–0.74% in canaries [46], 0.1–0.2% in zebra finches [47], and 0.2–0.4% in Bengalese finches (*Lonchura striata domestica*) [46–47]. Finding that neuronal addition rates remain modest even following reactive proliferation suggests that the primary functional output of the reactive proliferative event is not rapid neuronal replacement, but rather, the replenishment and remodeling of the glial environment in preparation for circuit regeneration in the subsequent breeding season.

Taken together, these results demonstrate that the dynamics and likely even the function of neuronal versus astrocytic turnover following environmentally-driven neural degeneration are fundamentally distinct. Several functional hypotheses have been proposed to explain the persistence of adult neurogenesis in the song circuit [19], though none has received unambiguous experimental support. One prominent hypothesis suggests that HVC neuronal addition enables adult song learning [43, 48–49]; however, this hypothesis is challenged by robust adult neurogenesis in species like the white-crowned sparrow that learn songs only as juveniles [19].

Alternative hypotheses posit that neuronal turnover facilitates the maintenance of previously learned songs through ongoing error correction and motor program updating, or replaces neurons damaged by the intense metabolic demands and potential excitotoxicity induced by prolonged high-rate singing during breeding season [19, 50–51]. Whatever the adaptive function of neuronal turnover in HVC, the parallel turnover of astrocytes likely serves to facilitate neuroprotection, debris clearance, and the re-establishment of a pro-regenerative glial environment following environmentally-triggered neuronal loss.

### Functional relevance of astrocyte turnover in environmentally-driven circuit remodeling

Adult astrocytes in songbirds likely serve multiple interconnected functions that support the remarkable seasonal plasticity of the song control circuit. Following telencephalic injury in zebra finches, reactive astroglia adopt a neuroprotective steroidogenic phenotype by up-regulating aromatase to produce local estradiol [35]. This localized up-regulation of estradiol synthesis suppresses secondary neuronal death, reduces cellular debris, modulates neuroinflammatory signaling and promotes injury-induced regeneration [52–56]. Although seasonally-induced neuronal death in the white-crowned sparrow is a natural, non-pathalogical phenomenon, it nonetheless drives local inflammatory-like responses [57] and likely elicits a similar neuroprotective phenotypic shift in the HVC astrocytic population for return to non-breeding homeostasis. To this end, mature astrocytes, and maybe even astrocytes generated during reactive proliferation, may be essential for orchestrating the return to non-breeding homeostasis and preparing the circuit for regeneration in the next breeding season.

Astrocyte turnover appears coordinated with neurogenesis: both total GFAP-positive and total Vimentin-numbers positive astrocytes in HVC positively correlate with total HuC/D neuron number (SI Table 4). This coordination is consistent with the well-established capacity of NPCs to generate both neurons and non-neuronal glia during both homeostatic [49,51,58–59] and seasonally-induced reactive proliferation (Figure 1D and SI Table 2), and with prior reports of astrocyte number scaling with neuron number in canaries [19]. The co-production of glia with neurons may provide both the physical scaffolding and the trophic microenvironment necessary for newly-born neurons to successfully migrate and integrate into their HVC targets [42, 60]. In rodents, astrocytes secrete neurotrophic factors including brain-derived neurotrophic factor (BDNF), vascular endothelial growth factor (VEGF), and insulin-like growth factor-1 (IGF-1) that promote neuronal survival, differentiation, and maturation [61–64]. Astrocytes also regulate neurotransmitter clearance, ion homeostasis, and metabolic substrate delivery to support functional incorporation of adult-born neurons [65–68], and participate directly in synaptogenesis through both physical tripartite synapse contacts and secretion of synaptogenic signals [69].

Given broad functional conservation of astrocyte biology across vertebrates ([70], but see [71]), including shared expression of core and function-specific astroglial markers, it is likely that many of these signaling and support mechanisms are conserved in the avian brain. Under this framework, the large-scale astrocyte turnover following environmentally-driven HVC degeneration likely serves both to clear the debris of dying neurons and to replenish the astrocytic population whose support functions are critical during the re-establishment of homeostasis and the subsequent regeneration of the song circuit and recovery of behavior.

### Identification of parenchymal progenitor cells beyond the canonical niche

In the adult mammalian parenchyma, astrocytes are overwhelmingly quiescent under homeostatic conditions, with proliferation negligible to absent outside the canonical neurogenic niches [72–73]. Following mild cortical ischemia, subsets of parenchymal astrocytes can transiently activate neurogenic programs [74–75], and *in vivo* astrocyte-to-neuron transition has been reported in mammals [76], but remains controversial [77]. The clearest evidence for parenchymal astrocytic proliferation comes from lineage-tracing studies in mouse cortex using multiple inducible Cre lines (e.g., SOX2-, GFAP-, and PDGFRα-CreER) combined with retroviral birth dating. These studies demonstrate that SOX2-positive astrocytes can produce migratory neuroblasts following mini-thrombotic lesions, with the subventricular zone contributing only a minor fraction of these cells [78]. Similar responses have been described in macaque cortex following ischemia, though definitive long-term neuronal survival has not been confirmed in primates [78]. Likewise, in songbirds, focal telencephalic injury triggers robust reactive astrogliosis around the site of the lesion and beyond the ventricular zone. This reactive astrogliosis is characterized by accumulation of aromatase-expressing reactive astrocytes [79]. More direct evidence for parenchymal astrocyte proliferation comes from studies of injured zebra finch hippocampus, where increased cell proliferation occurs within the lesion territory and alongside aromatase expression [36]. Together, these observations suggest that parenchymal glial cells across vertebrates retain latent proliferative capacity that can be unmasked by injury, though whether this capacity is engaged under non-injury, ecologically-relevant conditions had not previously been examined.

Despite these observations in mammals and birds, the general consensus has been that astrocyte neurogenic responses in the parenchyma follow a burst-and-fade pattern with very low overall efficiency and poor functional outcomes [80–81]. Most injury-induced neuroblasts die within days to weeks [80–81], with only an extremely small fraction cells maturing and functionally incorporating into existing circuits [81]. Prior to this study, whether parenchymal astrocytes proliferate and give rise to new progeny under non-injury homeostatic conditions – let alone during the recurring, environmentally-driven neural perturbation experienced by seasonally-breeding songbirds – remained unknown. Here, we report a population of SOX2-positive cells that are: 1) proliferative under homeostatic conditions, 2) capable of self-renewal, and, 3) likely contributing new fate-specified astrocytes and neurons within HVC. Critically, these cells expand their pool size in response to seasonally-driven, non-injury neuronal loss, establishing that parenchymal progenitor-like cells can respond to ecologically-relevant, physiological perturbation.

We identified SOX2-positive cells within HVC (Figures 5C and E) and the broader telencephalon, that incorporated EdU two hours prior to tissue harvesting (Figure 6A and B), confirming active proliferation under both homeostatic and degenerative conditions. Total number of SOX2-positive cell number within HVC increased significantly at two days following transition into nonbreeding conditions and retuned to baseline by seven days (Figure 6D and SI Table 3). The number of SOX2-positive cells incorporating EdU remained stable across the experimental time course (Figure 6C and SI Table 3), suggesting that the pool expansion is driven either strictly by vVZ NPCs, or that the expansion of pAPCs occurs in the day immediately preceding our earliest measurement point – a window not captured in the current study design. The absence of a significant correlation between vVZ NPC proliferation rates and pAPC proliferation rates supports the latter interpretation and suggests that pAPC pool expansion is a relatively autonomous response to the onset of seasonal neuronal loss rather than a downstream consequence of vVZ NPC activity.

Significant positive correlations were observed between EdU-incorporating SOX2-positive cells within HVC and: 1) the total population of SOX2-positive cells (Figure 6E and SI Table 4) consistent with self-renewal capacity and, 2) newly-born Vimentin-positive astrocytes (Figure 7A and SI Table 4), suggesting these proliferating cells give rise to fate-specified astrocytes. We find a significant correlation between newly-born HuC/D-expressing neurons and the total number of SOX2-positive cell number within HVC, but not with actively proliferating SOX2-positive cells (Figure 7C and SI Table 4). The absence of a correlation between proliferating SOX2-positive cells and neurons does not rule out the pAPCs as capable of giving rise to neurons. Rather, the lack of correlation could be a timing mismatch, wherein the bulk of new HVC neurons are born in the first day following onset of degeneration, prior to our two-day EdU labeling window. Alternatively, it is possible that the vVZ NPCs predominately give rise to the new neurons observed in HVC either: 1) at the same rate as SOX2 cells proliferating in HVC and self-renewing, or 2) in conjunction with some proportion of SOX2-positive cells.

The demonstration of homeostatic, progenitor-like astrocyte proliferation in adult brain parenchyma outside the canonical neurogenic niche fundamentally challenges the prevailing view that parenchymal astrocytes remain post-mitotic with negligible self-renewal capacity.

Importantly, these findings raise the possibility that parenchymal progenitor populations may represent a conserved but underappreciated substrate for neural resilience in species that face recurring, large-scale neural perturbations as a consequence of their ecological life histories. Rigorous confirmation that pAPCs are *bona fide* parenchymal neural progenitor cells will require experimental tools beyond those currently available in non-traditional model systems like the songbird, including: 1) inducible astrocyte-specific lineage tracing with controls excluding oligodendrocyte precursor cell-derived and subventricular zone migrant contributions; 2) *in vivo* proliferation analyses demonstrating sustained cell cycling; 3) functional fate readouts confirming persistent astrocyte replacement and neuronal maturation with synaptic integration; and 4) spatial transcriptomics validating a stable progenitor-like transcriptional state distinct from reactive astrogliosis.

### Overall impact and implications for neural resilience under environmental change

Our results demonstrate that both baseline astrocyte turnover rates and the proliferative capacity of parenchymal astrocytes in songbirds differ substantially from the mammalian pattern of negligible homeostatic proliferation and minimal parenchymal turnover outside canonical niches. These differences likely reflect evolutionary adaptations to life histories characterized by recurrent, large-scale, environmentally-driven neural remodeling. Such adaptations may have broad relevance for understanding how other seasonally-breeding or environmentally-sensitive vertebrates maintain neural circuit function and behavior across variable ecological conditions.

## ACKNOWLEDGEMENTS

We thank the Larson lab members for assistance with bird collections and care, K. Lent for assistance with bird collections, and D.J. Perkel and S.R. Larson for helpful discussions and advice. This work was supported by NIH R35 GM150562 to T.A.L and NSF NRT-ROL 2021791 to W.C.T..

## METHODS

### Animals

Eighty-three adult male Gambel’s white-crowned sparrows (*Zonotrichia leucophrys gambelli)* were collected a field site in Eastern Washington during the pre-breeding and post-breeding migrations. Minimum age was estimated by plumage coloration: birds with black and white crowns (adult; minimum 14 months) and birds with brown and white crowns (juveniles; minimum 2 months). All birds used in experiments had adult plumage and were at a minimum 14 months of age. Before the start of experiments, all birds were housed in group indoor aviaries in a short day photoperiod (SD; 8 hr light; 16 hr dark) for at least 10 weeks to ensure sensitivity to the stimulatory effects of long day photoperiod (LD) and testosterone (T) implants. All experimental protocols followed NIH, ALAAC, and USDA animal use guidelines and were approved by the University of Virginia Institutional Animal Care and Use Committee.

### Experimental Procedures

To test the timing and degree of integration of new cells into HVC following HVC degeneration and reactive proliferation, birds were transferred from SD to LD photoperiod and implanted with a 9 mm SILASTIC tube (DOW, 508-006, 1.47mm x 1.96mm) filled with crystalline testosterone (Sigma Aldrich, 1002747085), abbreviated LDT, to induce seasonal-like growth of the song circuit [17]. After 28 days of LDT, birds were transitioned to SD photoperiod and T implants were removed to induce degeneration of the song circuit (long day withdrawal; LDW) [17,45]. Beginning on the first day after transition into LDW, 5-bromo-2-deoxyuridine (BrdU; 50 mM in 15% DMSO and 0.7% saline at 50 mg/kg) was administered intramuscularly between Zeitgeber Time 4-6 (four to six hours after lights ON), for five consecutive days. Birds remained in LDW for varying durations as follows: LDT 28D (n=10), LDW 2D (n=8), LDW 4D (n=10), LDW 7D (n=8), LDW 10D (n=11), LDW 12D (n=9), LDW 14D (n=8), and LDW 28D (n=10). Two hours prior to tissue harvesting, birds were injected with 5-ethynyl-2′-deoxyuridine (EdU; 50 mM in 15% DMSO and 0.7% saline, 40.8 mg/kg, 12.3 mg/ml, Invitrogen) to label recently proliferative neural progenitor and stem cells.

### Song recording

All birds were housed in isolation in (IAC Acoustics) sound proofed chambers equipped with microphone recorders to collect song throughout the experiment. Syrinx software (J. Burt, www.syrinxpc.com) was used to compile and sort song recordings. To encourage singing, we provided birds with a mirror and played a 5-minute playback recording of Gambel’s white-crowned sparrow song every 15 minutes [17].

### Tissue collection and storage

Between Zeitgeber Time 4-6, birds were deeply anesthetized with isoflurane and brains were quickly removed. Within 2 minutes, brains were bisected at the midline and placed on dry ice for histology. Halved brains were sectioned at 40 um in the coronal plane with each section thaw-mounted serially in six sets. Tissue prior to and following sectioning was stored at −80C.

### Nissl staining

Every third section was Nissl stained using Thionin as follows. Tissue mounted on slides were fixed in 4% paraformaldehyde (in PBS) for 15 minutes, then rinsed with PBS (pH 7.4). Slides were then bathed in a 1.75 mM solution of thionin acetate in an aqueous solution of 0.57% sodium acetate, 0.0285% acetic acid, and water for up to 10 minutes. Slides were then dehydrated using increasing concentrations of ethanol: 70% ethanol for 1 minute, 95% ethanol for 30 seconds to several minutes, and 100% ethanol for 30 seconds. Slides were then dipped in xylenes and mounted with DPX mounting medium (Electron Microscopy Sciences; Cat. No 13512). Our full detailed protocol for Nissl staining is available here: https://dx.doi.org/10.17504/protocols.io.261ger8edl47/v1.

### Immunohistochemical Staining

Tissue preparation for all immunohistochemistry was performed as follows. Slides were slowly defrosted at room temperature and then fixed in 4% paraformaldehyde (PFA in phosphate buffered saline (PBS); (10X stock: 80g NaCl, 2g KCl, 14.4g Na_2_PO_4_, 2.4g H_2_KPO_4_, 1L water; diluted to 1X; pH 7.4) for 15 minutes. Slides were with either PBS (BrdU/astrocyte markers, EdU/SOX2) or PBS with 0.5% Triton X and 0.5% DMSO (PDTX; BrdU/neuron markers).

To visualize the BrdU antigen, fixed tissue was incubated in 100% methanol (MeOH) for 15 minutes at −20C, then gradually rehydrated with an additional 25% PBS concentration every 3 minutes. After a final rinse in 100% PBS for 15 min, slides were briefly rinsed in distilled water and incubated in freshly made 2N HCl for 30 minutes at 37C. Slides were then incubated in a 0.03% hydrogen peroxide and subsequently an avidin/biotin kit solution (Vector Laboratories), both in PBS, for blocking of endogenous avidin and biotin and quenching of endogenous peroxidases. Slides were rinsed three times in either PBS (BrdU/astrocyte markers) or PDTX (BrdU/neuron markers) for 15 minutes each before blocking with 5% heat inactivated goat serum (Vector Laboratories; SKU S-1000-20 in PBS). BrdU was labeled by incubating tissue with the mouse-anti-BrdU antibody (Developmental Studies Hybridoma Bank, G3G4; 1:350 in 5% heat inactivated goat serum) overnight at 4C or at room temperature for 2 hours. Tissue was rinsed four times in PBS or PDTX for 15 minutes each. A goat-anti-mouse biotinylated IgG secondary antibody (Vector Laboratories; SKU BA-9200-1.5; 1:200 in % heat inactivated goat serum) was incubated for 2 hours at room temperature followed by amplification with the avidin-biotin peroxidase (ABC) complex (Vector Laboratories; SKU PK-6100) as per manufacturer’s protocol. Visualization of the secondary antibody was obtained by treating tissue with a nickel DAB stain (0.05% DAB, 0.05% nickel ammonium sulfate, 0.015% hydrogen peroxide in PBS or TBS, pH 7.0-7.5) for a dark purple to black stain to label DNA containing BrdU.

To label new and mature astrocytes and neurons, BrdU-labeled slides were then labeled for either: immature astrocytes targeting vimentin (mouse anti-vimentin; Invitrgoen clone V9; 1:200 in 5% heat-inactivated goat serum) [30]; proliferative and mature astrocytes targeting GFAP (rabbit-anti-GFAP, DAKO; 1:200 [30]); or mature neurons targeting HuC/D (mouse-anti HuC/HuD, Invitrogen A21271; 1:100 in 5% heat-inactivated goat serum [17]). After rinses as described above, slides were incubated in goat anti-mouse or goat anti-rabbit biotinylated IgG secondary antibody (1:200 in 5% heat inactivated goat serum); amplified with the Vector ABC complex kit as above; and detected with DAB stain without nickel (0.05% DAB, 0.015% hydrogen peroxide in PBS or TBS, pH 7.0-7.5) for a brown stain in astrocyte or neuron marker positive cells. After labeling of BrdU and lineage-specific markers in the same tissue, slides were then dehydrated using increasing concentrations of ethanol: 70% ethanol for 1 minute, 95% ethanol for 30 seconds to several minutes, and 100% ethanol for 30 seconds. Slides were then dipped in xylenes and mounted with DPX mounting medium (Electron Microscopy Sciences; Cat. No 13512).

To visualize recent proliferation of neural progenitor-like cells, tissue was incubated in 15% methanol for 15 minutes at −20C. Slides were then incubated in a 0.03% hydrogen peroxide and subsequently an avidin/biotin kit solution (Vector Laboratories) for blocking of endogenous avidin and biotin and quenching of endogenous peroxidases, both in PBS. Developing solution from the EdU Click-it Kit (Thermo Fisher; Ref No. C10269; biotin azide antibody at 1:800) (component amounts published in online protocol) was then made and applied to slides which were incubated for 2-3 hours at room temperature or overnight at 4C. Solution was then thoroughly rinsed and avidin-biotin peroxidase (ABC) complex (Vector Laboratories; SKU PK-6100) was added for signal amplification as per manufacturer’s protocol. After thorough rinsing in PBS, DAB staining solution (0.05% DAB, 0.05% nickel ammonium sulfate, 0.015% hydrogen peroxide in PBS or TBS, pH 7.0-7.5) was added to the slides to induce a dark purple to black stain in EdU positive cells. To ensure quenching of staining reaction and prevent excessive background, slides were incubated in a 1% ethylenediaminetetraacetic acid (EDTA) solution.

After multiple rinses in PBS, rabbit-anti-SOX2 (Abcam; Cat No. AB97959; 1:200 in 5% heat-inactivated goat serum) was added to slides and incubated for about 2 hours at room temperature or overnight at 4C. Slides were then thoroughly rinsed in PBS and incubated with a biotinylated IgG goat anti-rabbit secondary antibody (Vector; BA-1000-1.5; 1:200 in 5% heat inactivated goat serum). Slides were thoroughly rinsed in PBS and avidin-biotin peroxidase (ABC) complex (Vector Laboratories; SKU PK-6100) was added for signal amplification as per manufacturer’s protocol. Slides were then thoroughly rinsed and DAB staining solution without nickel ammonium sulfate (0.05% DAB, 0.015% hydrogen peroxide in PBS or TBS, pH 7.0-7.5). After EdU/SOX2 labeling was completed, slides were then dehydrated using increasing concentrations of ethanol: 70% ethanol for 1 minute, 95% ethanol for 30 seconds to several minutes, and 100% ethanol for 30 seconds. Slides were then dipped in xylenes and mounted with DPX mounting medium (Electron Microscopy Sciences; Cat. No 13512).

Detailed immunostaining protocols are available here: BrdU/vimentin, https://dx.doi.org/10.17504/protocols.io.5qpvoe47bl4o/v1; BrdU/GFAP, https://dx.doi.org/10.17504/protocols.io.261ge193jv47/v1; BrdU/Hu, https://dx.doi.org/10.17504/protocols.io.14egn1r8zv5d/v1; SOX2/EdU, https://dx.doi.org/10.17504/protocols.io.q26g77nb1gwz/v1.

### Immunofluorescent Staining

Fluorescent staining was performed with selected markers. Tissue was permeabilized with 15% methanol for 15 minutes at −20C. For EdU/SOX2-labeled tissue, developing solution made with the EdU Click-it Kit (Thermo Fisher; Ref No. C10269; Alexa Fluor 561-conjugated biotin azide antibody at 1:800) (component amounts published in online protocol) was incubated with tissue for 2-3 hours at room temperature or at 4C overnight in the dark. Tissue was then thoroughly rinsed and rabbit-anti-SOX2 (Abcam; Cat No. AB97959; 1:200 in 5% heat-inactivated goat serum) was applied and incubated for around 2 hours at room temperature or overnight at 4C in the dark. Tissue was then thoroughly rinsed and goat anti-rabbit IgG Alexa Fluor 488 secondary antibody (Invitrogen; 1:200 in 5% heat inactivated goat serum) was applied and incubated for around 2 hours at room temperature or at 4C overnight in the dark. Tissue was thoroughly rinsed and mounted in Fluoromount-G™ with DAPI (Invitrogen). For SOX2/vimentin labeled tissue, SOX2 labeling was performed as stated above. After SOX2 labeling, tissue was thoroughly rinsed and mouse anti-vimentin (Invitrogen; clone V9; Ref No. 14-9897-82; 1:200 in 5% heat-inactivated goat serum) was applied and incubated for around 2 hours at room temperature or 4C overnight in the dark. Tissue was then thoroughly rinsed and an Alexa Fluor 647 conjugated IgG goat anti-mouse secondary antibody (Invitrogen, 1:200 in 5% heat-inactivated goat serum). Tissue was thoroughly rinsed and mounted in Fluoromount-G™ with DAPI.

### Image Processing

Representative images of chromogen stained tissue were captured with a high-resolution slide imager (Huron Digital Pathology; TissueScope™ LE120). Specific magnifications used for each image are highlighted in figures and figure captions. Representative images of immunofluroescently-labeled tissue were captured on a Nikon ECLIPSE Ti2 inverted microscope. Specific objective lens magnifications used for each image are highlighted in figures and figure captions. False coloring of black and white images was performed in Photoshop.

### Song analysis

To measure the song percent complete, we randomly selected 30 songs from time points of interest. For each individual song, we counted the number of syllables out of five that the bird sang, and then divided syllables observed by syllables expected to obtain a percentage sang out of the complete song. To measure song rate, we counted the total number of songs sang (of any length) on the last full day prior to the day of sacrifice to obtain songs per day. Averages and standard error for each time point were compiled and data was analyzed using a one-way ANOVA (JMP Pro, version 18).

### Morphometric Measurements

One hemisphere was randomly selected for Nissl staining to obtain morphometric measurements of HVC and RA. Volumes were determined by tracing outlines of these nuclei on paper using a microprojector (Bausch-Lomb). Tracings were scanned and analyzed using ImageJ (v1.46, NIH; http://rsb.info.nih.gov/ij/). Areas were calculated and total volume was obtained by using the formula for a truncated cone [17]. Mean and SEM for each experimental time point were then compiled and data was analyzed using a one-way ANOVA (JMP Pro, version 18).

### Immunohistochemistry Quantification

All astrocyte-marker labeled tissue was double labeled with an astrocyte marker (vimentin or GFAP) and BrdU, and a subset of 3 HVCs were counted from each brain for single and double labeled cells within HVC. The same counting protocol was applied to EdU/SOX2 labeled tissue in HVC.

As with the astrocyte markers, HuC/D was double labeled with BrdU, and a subset of 3 sections with HVC were counted for single and double labeled cells of each marker. VZ counts were performed in tissue labeled for EdU and SOX2. For the astrocyte and neuron markers, sections were counted using the inbuilt grid on a Zeiss Axioplan 2 microscope at 400x. Cell density for each marker was calculated by multiplying the raw cell count by the grid field of view, and then the counts for each of the three sections were averaged to obtain a mean. Cell number was calculated by multiplying the observed HVC volume by the mean cell density for each individual brain. For EdU/SOX2 labeled tissue, slides were imaged at high resolution using a slide imager (Huron Digital Pathology; TissueScope™ LE120) and the number of EdU positive cells labeled in the ventral VZ were counted from the hillock of the VZ medial to HVC, out to the most lateral extension of the vVZ in all sections that exhibited arching of the VZ [17]. For each marker, means and SEMs were compiled for each time point and data was analyzed using a one-way ANOVA (JMP Pro, version 18). All counts reported here are unilateral; however, the hemisphere used for each marker was randomized.

### Regression Analysis

Linear regressions were performed using Pearson’s correlation with specified variables using (JMP Pro, version 18). Birds for which data was missing for the variable of interest were removed from the dataset and not included in the respective analysis.

## Author Contributions

Conceptualization: WCT, JMB, TAL

Data curation: WCT, PEC, JMB, TAL

Formal analysis: WCT, TAL

Funding acquisition: WCT, TAL

Investigation: WCT, SLS, PEC, AM, JMB

Methodology: WCT, SLS, PEC, JMB

Project administration: TAL

Resources: TAL

Supervision: TAL

Validation: WCT, TAL

Visualization: WCT, TAL

Writing: WCT, TAL

