## Supplementary material for "Parenchymal precursor cells act in concert with canonical neural progenitors to facilitate neural circuit recovery following neuronal loss": SI Table 1

**SI Table 1.** Confirmation of degradation and reactive proliferation

| | LDT | 2D | 4D | 7D | 10D | 12D | 14D | 28D | ANOVA |
|---|---|---|---|---|---|---|---|---|---|
| HVC Volume ($mm^3$) | - | 0.465 ± 0.03 | 0.453 ± 0.02 | 0.394 ± 0.03 | 0.436 ± 0.02 | 0.359 ± 0.04 | 0.407 ± 0.03 | 0.393 ± 0.03 | $F_{(6,57)}=1.6894$ $p=0.1402$ |
| Neuron Number ($x10^3$) | - | 33 ± 2 | 34 ± 2 | 30 ± 3 | 32 ± 3 | 26 ± 4 | 29 ± 3 | 30 ± 2 | $F_{(6,48)}=1.0290$ $p=0.4182$ |
| EdU Positive Cells in vVZ | - | 1254 ± 165[a,b] | 1989 ± 569[a] | 567 ± 99[b] | 1102 ± 168[a,b] | 863 ± 237[a,b] | 699 ± 158[b] | 1043 ± 169[a,b] | **$F_{(6,48)}=3.1368$ $p=0.0113$** |
| Song Percent Complete | 80.23 ± 1.87[a] | 28.05 ± 11.02[b] | 39.55 ± 18.87[b] | 43.30 ± 16.11[b] | 6.19 ± 6.19[b] | 28.36 ± 15.07[b] | 15.00 ± 15.00[b] | 18.38 ± 12.27[b] | **$F_{(7,110)}=18.4631$ $p<0.0001$** |

All values are mean ± S.E.M. Superscript letters denote significant differences across treatment groups with post-hoc Tukey tests.
