## Supplementary material for "Parenchymal precursor cells act in concert with canonical neural progenitors to facilitate neural circuit recovery following neuronal loss": SI Table 2

**SI Table 2.** Summary of reactive proliferation progeny

| | 2D | 4D | 7D | 10D | 12D | 14D | 28D | ANOVA |
|---|---|---|---|---|---|---|---|---|
| **HVC Cell Number** | | | | | | | | |
| Total HuC/D* | 33 ± 2 | 34 ± 2 | 30 ± 3 | 32 ± 3 | 26 ± 4 | 29 ± 3 | 30 ± 2 | $F_{(6,48)}$=1.0290 p=0.4182 |
| HuC/D & BrdU (per day BrdU injection) | 1157 ± 111[a] | 537 ± 70[b] | 328 ± 51[b] | 318 ± 44[b] | 367 ± 74[b] | 344 ± 44[b] | 364 ± 57[b] | **$F_{(6,48)}$=19.6345 p<0.0001** |
| HuC/D & BrdU (unadjusted) | 2316±222 | 2149±280 | 1641±256 | 1591±221 | 1709±364 | 1752±221 | 1727±256 | $F_{(6,48)}$=1.0801 p=0.3877 |
| Total Vimentin | 34874 ± 5048 | 29335 ± 4185 | 24954 ± 4529 | 27235 ± 1288 | 23388 ± 2856 | 26442 ± 3668 | 28294 ± 4282 | $F_{(6,46)}$=0.8511 p=0.5377 |
| Vimentin & BrdU (per day of BrdU injection) | 5055 ± 584[a] | 1961 ± 252[b] | 1261 ± 219[b] | 1671 ± 201[b] | 1767 ± 319[b] | 1590 ± 228[b] | 2019 ± 506[b] | **$F_{(6,46)}$=12.3438 p<0.0001** |
| Vimentin & BrdU (unadjusted) | 10109 ± 1167 | 7845 ± 1008 | 6308 ± 1096 | 8357 ± 1005 | 8836 ± 1597 | 7953 ± 1142 | 10097 ± 2532 | $F_{(6,46)}$=0.8050 p=0.5713 |
| **Ratios** | | | | | | | | |
| BrdU & HuC/D to BrdU & Vimentin | 0.2584 ± 0.0669 | 0.3108 ± 0.0454 | 0.3398 ± 0.0982 | 0.1951 ± 0.0297 | 0.1785 ± 0.0195 | 0.2344 ± 0.0281 | 0.2143 ± 0.0780 | $F_{(6,37)}$=1.1241 p=0.3675 |
| BrdU & HuC/D Day X to Day 2 LDW | - | 0.4644 ± 0.0604 | 0.2836 ± 0.0442 | 0.2644 ± 0.0434 | 0.3168 ± 0.0638 | 0.2977 ± 0.0382 | 0.3117 ± 0.0648 | $F_{(5,39)}$=1.886 p=0.1190 |
| BrdU & HuC/D to total HuC/D | 0.0395 ± 0.0061[a] | 0.0157 ± 0.0011[b] | 0.0115 ± 0.0022[b] | 0.0093 ± 0.0013[b] | 0.0133 ± 0.0014[b] | 0.0125 ± 0.0018[b] | 0.0123 ± 0.0019[b] | **$F_{(6,46)}$=13.8425 p<0.0001** |
| BrdU & Vimentin Day X to Day 2 LDW | - | 0.3880 ± 0.0499 | 0.2496 ± 0.0434 | 0.3306 ± 0.0398 | 0.3496 ± 0.0632 | 0.3147 ± 0.0452 | 0.4329 ± 0.1117 | F(5,40)=1.0258 p=0.4156 |
| BrdU & Vimentin to total Vimentin | 0.1460 ± 0.0208[a] | 0.0693 ± 0.0068[b] | 0.0530 ± 0.0054[b] | 0.0608 ± 0.0062[b] | 0.0747 ± 0.0084[b] | 0.0704 ± 0.0174[b] | 0.0684 ± 0.0129[b] | **F(6,46)=6.1953 p<0.0001** |

*Same data in SI Table 1, here for ease of comparisons

All values are mean ± S.E.M. Superscript letters denote significant differences across treatment groups with post-hoc Tukey tests.
