## Supplementary material for "Parenchymal precursor cells act in concert with canonical neural progenitors to facilitate neural circuit recovery following neuronal loss": SI Table 3

**SI Table 3.** Verification of progenitor-like capacity of pAPCs

| | 2D | 4D | 7D | 10D | 12D | 14D | 28D | ANOVA |
|---|---|---|---|---|---|---|---|---|
| **HVC Cell Number** | | | | | | | | |
| Total GFAP | 30015 ± 3021 | 30412 ± 3309 | 30974 ± 3377 | 28874 ± 3431 | 27866 ± 2803 | 32318 ± 3548 | 29806 ± 4280 | $F_{(6,47)}$=0.1754 p=0.9822 |
| GFAP & BrdU (per day of BrdU injection) | 5440 ± 626[a] | 1806 ± 371[b] | 1780 ± 372[b] | 1925 ± 447[b] | 2037 ± 369[b] | 1489 ± 195[b] | 1434 ± 300[b] | **$F_{(6,47)}$=12.0969 p=<0.0001** |
| GFAP & BrdU (unadjusted) | 10879 ± 1253 | 7225 ± 1485 | 8901 ± 1862 | 9623 ± 2235 | 10186 ± 1844 | 7446±974 | 7175±1500 | $F_{(6,47)}$=0.8902 p=0.5098 |
| Total Sox2 | 26925 ± 3296[a] | 18544 ± 3120[a,b] | 13684 ± 2311[b] | 15645 ± 2302[a,b] | 12917 ± 2496[b] | 15265 ± 3041[a,b] | 19359 ± 2876[a,b] | **$F_{(6,45)}$=2.7007 p=0.0252** |
| Sox2 & EdU (2hr pulse) | 1891 ± 211 | 1836 ± 373 | 1741 ± 293 | 1628 ± 337 | 1702 ± 491 | 1928 ± 185 | 1377 ± 232 | $F_{(6,45)}$=0.3355 p=0.9146 |
| **Ratios** | | | | | | | | |
| BrdU & GFAP Day X to Day 2 LDW | - | 0.3321±0.0682 | 0.3273±0.0685 | 0.3538±0.0822 | 0.3745±0.0678 | 0.2738±0.0358 | 0.2751±0.0623 | $F_{(5,45)}$=0.3926 p=0.8509 |
| BrdU & GFAP to total GFAP | 0.1910±0.0236[a] | 0.0582 ± 0.0087[b] | 0.0605 ± 0.0112[b] | 0.0674 ± 0.0139[b] | 0.0730 ± 0.0109[b] | 0.0510 ± 0.0092[b] | 0.0548 ± 0.0109[b] | **$F_{(6,46)}$=13.7143 p<0.0001** |

All values are mean ± S.E.M. Superscript letters denote significant differences across treatment groups with post-hoc Tukey tests.
