## Supplementary material for "Parenchymal precursor cells act in concert with canonical neural progenitors to facilitate neural circuit recovery following neuronal loss": SI Table 4

**SI Table 4.** Pearson's Correlations between various cell metrics

| **Correlation Variables** | $r^2$ | **p-value** |
|---|---|---|
| *Relationship astrocytes and neurons* | | |
| Total GFAP to Total Hu | 0.3218 | **<0.0001** |
| Total Vimentin to Total Hu | 0.2536 | **0.0002** |
| BrdU/Vimentin to BrdU/Hu | 0.4645 | **<0.0001** |
| *Progeny of vVZ NPCs* | | |
| EdU in vVZ to BrdU/Vimentin/pulse | 0.0012 | 0.8150 |
| EdU in vVZ to BrdU/Hu/pulse | 0.0201 | 0.3216 |
| EdU in vvZ to Total Sox2 | 0.0037 | 0.6775 |
| EdU in vvZ to EdU/Sox2 | 0.0187 | 0.3438 |
| EdU in vVZ to total Vimentin | 0.0003 | 0.6935 |
| EdU in vVZ to total Hu | 0.0485 | 0.1243 |
| *Self-renewal pAPCs* | | |
| Total Sox2 to Sox2/EdU | 0.2325 | **<0.0001** |
| Total Sox2 to Total GFAP | 0.0638 | 0.0769 |
| Total Sox2 to BrdU/GFAP/pulse | 0.2798 | **<0.0001** |
| Sox2/EdU to BrdU/GFAP | 0.0404 | 0.1614 |
| *Progeny of pAPCs* | | |
| EdU/Sox2 to BrdU/Vimentin/pulse | 0.0922 | **0.0358** |
| Total Sox2 to BrdU/Vimentin/pulse | 0.3282 | **<0.0001** |
| EdU/Sox2 to BrdU/Hu/pulse | 0.0002 | 0.9270 |
| Total Sox2 to BrdU/Hu/pulse | 0.1721 | **0.0020** |
