## Supplementary material for "Parenchymal precursor cells act in concert with canonical neural progenitors to facilitate neural circuit recovery following neuronal loss": SI Table 5

**SI Table 5.** Raw song percent complete data from individual birds

*If no number, bird sang less than 20 songs and data not included. Zeros represent birds with no songs the given day.

| BirdID | Condition | % Complete LDT27 | % Complete LDW |
|---|---|---|---|
| 3612 | 10D | 92.50 | 0.00 |
| 3541 | 10D | 80.63 | 0.00 |
| 3790 | 10D | 73.75 | 0.00 |
| 3742 | 10D | 49.38 | 0.00 |
| G0087 | 10D | 97.50 | 0.00 |
| G0093 | 10D | 64.38 | 43.33 |
| G0084 | 10D | | |
| G0162 | 10D | 90.59 | 0.00 |
| G0130 | 12D | 81.88 | |
| G0014 | 12D | 95.00 | 0.00 |
| G0029 | 12D | 69.38 | 0.00 |
| G0044 | 12D | 72.50 | 100.00 |
| G0050 | 12D | 68.75 | |
| G0012 | 12D | 91.88 | 0.00 |
| G0131 | 12D | 55.00 | 0.00 |
| G0193 | 12D | 96.88 | 0.00 |
| G0176 | 12D | 88.75 | 86.88 |
| G0175 | 12D | 80.00 | 40.00 |
| 3704 | 14D | 85.63 | 0.00 |
| 3719 | 14D | 93.13 | 0.00 |
| 3844 | 14D | 96.15 | |
| 3682 | 14D | 87.90 | 0.00 |
| G0053 | 14D | 74.38 | |
| G0023 | 14D | 85.81 | |
| G0174 | 14D | 84.38 | 75.00 |
| G0178 | 14D | 88.75 | 0.00 |
| G0201 | 14D | 100.00 | |
| 3654 | 28D | 63.75 | 96.88 |
| 3706 | 28D | 78.75 | 0.00 |
| 3728 | 28D | 70.00 | 0.00 |
| 3593 | 28D | 73.75 | 0.00 |
| 3808 | 28D | 87.50 | 0.00 |
| 3492 | 28D | 96.25 | 86.88 |
| G0075 | 28D | 69.70 | 0.00 |
| G0115 | 28D | 97.50 | 0.00 |
| G0060 | 28D | 99.39 | 0.00 |
| G0170 | 28D | 96.00 | 0.00 |
| 3402 | 2D | 90.63 | 0.00 |
| 3744 | 2D | 57.50 | 0.00 |
| 3590 | 2D | 58.13 | 57.50 |
| 3691 | 2D | 96.25 | 73.13 |
| G0006 | 2D | 94.38 | 0.00 |
| G0118 | 2D | 100.00 | |
| G0133 | 2D | 89.38 | |
| 3426 | 2D | 43.75 | 41.88 |
| G0136 | 2D | 89.38 | 51.88 |
| G0192 | 2D | 92.50 | 0.00 |
| 3683 | 4D | 71.25 | |
| 3722 | 4D | 97.50 | |
| G0026 | 4D | 76.88 | 100.00 |
| G0022 | 4D | 93.75 | 0.00 |
| G0045 | 4D | | |
| G0094 | 4D | | 76.88 |
| G0030 | 4D | 40.00 | 0.00 |
| G0031 | 4D | 87.50 | 100.00 |
| G0186 | 4D | 94.38 | 0.00 |
| G0191 | 4D | | 0.00 |
| 3665 | 7D | 53.75 | 55.00 |
| 3599 | 7D | 74.38 | 0.00 |
| 3746 | 7D | 95.63 | 99.38 |
| 3632 | 7D | 76.25 | 0.00 |
| G0016 | 7D | 85.63 | 68.75 |
| G0069 | 7D | 88.75 | |
| G0055 | 7D | 61.67 | |
| G0172 | 7D | 57.58 | 80.00 |
| G0179 | 7D | 58.75 | 0.00 |
| 3620 | LDT | 66.25 | 0.00 |
| 3709 | LDT | 80.00 | 0.00 |
| G0101 | LDT | 73.13 | 0.00 |
| G0112 | LDT | 73.14 | 0.00 |
| G0073 | LDT | weird syllables excluded | |
| G0106 | LDT | weird syllables, extra syllables | |
| G0135 | LDT | weird syllables, extra syllables | |
