## Supplementary material for "Parenchymal precursor cells act in concert with canonical neural progenitors to facilitate neural circuit recovery following neuronal loss": SI Table 6

**SI Table 6.** Raw morphometric and cytoarchitecture data from individual birds

| Bird ID | Time Point | Volume HVC | Hu+ Density | Hu+ Number | BrdU+/Hu+ per day Density | BrdU+/Hu- per day Density | BrdU+/Hu+ Density | BrdU+/Hu- Density | BrdU+/Hu+ per day Number | BrdU+/Hu- per day Number | BrdU+/Hu+ Number | BrdU+/Hu- Number | EdU+/Sox2- Density | Sox2+/EdU- Density | Sox2+/EdU + Density | EdU+/Sox2- Number | Sox2+/EdU- Number | Sox2+/EdU + Number | Sox2 Total | Vim+ Density | GFAP+ Density | Vim+ Number | GFAP+ Number | Vim+/BrdU+ density | GFAP+/BrdU+ density | Vim+/BrdU+ number | GFAP+/BrdU+ number | Vim+/BrdU+ density per day | GFAP+/BrdU+ density per day | Vim+/BrdU+ number per day | GFAP+/BrdU+ number per day | vVZ EdU+ Number |
|---|---|---|---|---|---|---|---|---|---|---|---|---|---|---|---|---|---|---|---|---|---|---|---|---|---|---|---|---|---|---|---|---|
| 3541 | 10D | 0.4110 | 51600 | 21208 | 560 | 1360 | 2800 | 6800 | 230 | 559 | 1151 | 2795 | | | | | | | | 54400 | 81400 | 22358 | 33455 | 13600 | 51600 | 5590 | 21208 | 2720 | 10320 | 1118 | 4242 | 1013 |
| 3612 | 10D | 0.4886 | 75200 | 36743 | 560 | 880 | 2800 | 4400 | 274 | 430 | 1368 | 2150 | 6000 | 40132 | 2132 | 2391 | 19608 | 1042 | 20650 | 64400 | 38800 | 31466 | 18958 | 14600 | 20000 | 7134 | 9772 | 2920 | 4000 | 1427 | 1954 | 1548 |
| 3742 | 10D | 0.3824 | 88800 | 33957 | 720 | 1546 | 3600 | 7732 | 275 | 591 | 1377 | 2957 | 4600 | 27400 | 2000 | 1759 | 10478 | 765 | 11243 | 60800 | 54000 | 23250 | 20650 | 11600 | 6000 | 4436 | 2294 | 2320 | 1200 | 887 | 459 | 1344 |
| 3790 | 10D | 0.4089 | | | 640 | 800 | 3200 | 4000 | 262 | 327 | 1308 | 1636 | 6400 | 25600 | 5600 | 2617 | 10468 | 2290 | 12758 | 77068 | 74668 | 31513 | 30532 | 26800 | 16932 | 10959 | 6923 | 5360 | 3386 | 2192 | 1385 | 504 |
| G0027 | 10D | 0.5013 | 73200 | 36695 | 480 | 840 | 2400 | 4200 | 241 | 421 | 1203 | 2105 | 4400 | 38200 | 3200 | 2206 | 19150 | 1604 | 20754 | 59400 | 76800 | 29777 | 38500 | 20800 | 15600 | 10427 | 7820 | 4160 | 3120 | 2085 | 1564 | 1464 |
| G0084 | 10D | 0.4860 | 84800 | 41213 | 1140 | 1200 | 5700 | 6000 | 554 | 583 | 2770 | 2916 | 10400 | 36000 | 6000 | 5054 | 17496 | 2916 | 20412 | 59000 | 82800 | 28674 | 40241 | 25034 | 24800 | 12167 | 12053 | 5007 | 4960 | 2433 | 2411 | 900 |
| G0087 | 10D | | 77200 | | 1240 | 1866 | 6200 | 9332 | | | | | 5800 | 28200 | 3600 | | | | | 58800 | 52400 | | | 21400 | 7600 | | | 4280 | 1520 | | | 1649 |
| G0093 | 10D | 0.4868 | | | | | | | | | | | | | | | | | | 54268 | | 26418 | | 12932 | | 6295 | | 2586 | | 1259 | | |
| G0162 | 10D | 0.3197 | 73600 | 23530 | 1226 | 1466 | 6132 | 7332 | | 469 | 1960 | 2344 | 6400 | 21600 | 3600 | 2046 | 6906 | 1151 | 8056 | 76400 | 61868 | 24425 | 19779 | 30800 | 22800 | 9847 | 7289 | 6160 | 4560 | 1969 | 1458 | 396 |
| G0012 | 12D | 0.4220 | 82800 | 34942 | 800 | 1120 | 4000 | 5600 | 338 | 473 | 2318 | 3245 | 2800 | 40400 | 3732 | 1182 | 17049 | 1575 | 18624 | 61200 | 86000 | 25826 | 36292 | 19800 | 20400 | 8356 | 8609 | 3960 | 4080 | 1671 | 1722 | 690 |
| G0014 | 12D | 0.3718 | 62800 | 23349 | 906 | 1174 | 4532 | 5868 | 337 | 436 | 1685 | 2182 | | | | | | | | 72932 | 87000 | 27116 | 32347 | 19068 | 21000 | 7089 | 7808 | 3814 | 4200 | 1418 | 1562 | |
| G0029 | 12D | 0.3013 | | | | | | | | | | | 8800 | 11200 | 1200 | 2651 | 3375 | 362 | 3736 | 66000 | 86000 | 19886 | 25912 | 19600 | 11600 | 5905 | 3495 | 3920 | 2320 | 1181 | 699 | |
| G0044 | 12D | 0.2824 | 75600 | 21349 | 680 | 1040 | 3400 | 5200 | 192 | 294 | 960 | 1468 | 4400 | 33600 | 7600 | 1243 | 9489 | 2146 | 11635 | 59600 | 77468 | 16831 | 21877 | 31600 | 41200 | 8924 | 11635 | 6320 | 8240 | 1785 | 2327 | 1119 |
| G0050 | 12D | 0.4381 | 77733 | 34055 | 2667 | 15067 | 2667 | 15067 | 533 | 6601 | 1168 | 6601 | 8000 | 29600 | 4800 | 3505 | 12968 | 2103 | 15071 | 58400 | 74268 | 25585 | 32537 | 32400 | 37600 | 14194 | 16473 | 6480 | 7520 | 2839 | 3295 | 567 |
| G0130 | 12D | 0.1438 | 65200 | 9376 | 680 | 1280 | 3400 | 6400 | 98 | 184 | 489 | 920 | 8000 | 19800 | 2800 | 1150 | 2847 | 403 | 3250 | 76200 | 80600 | 10958 | 11590 | 25000 | 25200 | 3595 | 3624 | 5000 | 5040 | 719 | 725 | 825 |
| G0131 | 12D | | 59000 | | 880 | 5960 | 4400 | 29800 | | | | | 10000 | 33800 | 6800 | | | | | 67600 | 88000 | | | 29600 | 42200 | | | 5920 | 8440 | | | 2130 |
| G0175 | 12D | 0.3554 | | | | | | | | | | | 4280 | 51760 | 2920 | 1521 | 18396 | 1038 | 19433 | 63650 | | 22621 | | 16100 | | 5722 | | 3220 | | 1144 | | 510 |
| G0176 | 12D | 0.4201 | 60533 | 25430 | 933 | 7173 | 4667 | 35867 | 392 | 3014 | 1961 | 15068 | 5600 | 40800 | 11200 | 2353 | 17140 | 4705 | 21845 | | 73200 | | 30751 | | 30800 | | 12939 | | 6160 | | 2588 | |
| G0193 | 12D | 0.4972 | 72800 | 36196 | 1360 | 2080 | 6800 | 10400 | 676 | 1034 | 3381 | 5171 | 5600 | 17000 | 2600 | 2784 | 8452 | 1293 | 9745 | 77000 | 63600 | 38284 | 31622 | 34000 | 34000 | 16905 | 16905 | 6800 | 6800 | 3381 | 3381 | 198 |
| 3682 | 14D | 0.5294 | 80668 | 42706 | 520 | 960 | 2600 | 4800 | 275 | 508 | 1376 | 2541 | 13200 | 28000 | 3200 | 6988 | 14823 | 1694 | 16517 | 82000 | 84800 | 43411 | 44893 | 13200 | 6932 | 6988 | 3670 | 2640 | 1386 | 1398 | 734 | |
| 3704 | 14D | 0.5418 | 75200 | 40743 | 680 | 720 | 3400 | 3600 | 368 | 390 | 1842 | 1950 | 11800 | 62400 | 3600 | 6393 | 33808 | 1950 | 35759 | 28800 | 58800 | 15604 | 31858 | 26800 | 8800 | 14520 | 4768 | 5360 | 1760 | 2904 | 954 | 459 |
| 3719 | 14D | 0.4179 | 61200 | 25575 | 680 | 1240 | 3400 | 6200 | 284 | 518 | 1421 | 2591 | 5600 | 42400 | 6400 | 2340 | 17719 | 2675 | 20394 | 53600 | 65600 | 22399 | 27414 | 12800 | 20400 | 5349 | 8525 | 2560 | 4080 | 1070 | 1705 | 705 |
| 3844 | 14D | 0.4474 | 65732 | 29408 | 960 | 586 | 4800 | 2932 | 429 | 262 | 2148 | 1312 | 7200 | 35200 | 4800 | 3221 | 15748 | 2148 | 17896 | | | | | | | | | | | | | 1350 |
| G0181 | 14D | 0.2450 | 74400 | 18228 | | | 5800 | 9800 | | | 1421 | 2401 | 12400 | 20000 | 4000 | 3038 | 4900 | 980 | 5880 | | | | | | | | | | | | | |
| G0023 | 14D | 0.4243 | 76800 | 32586 | 920 | 2000 | 4600 | 10000 | 390 | 849 | 1952 | 4243 | 9000 | 15600 | 3600 | 3819 | 6619 | 1527 | 8147 | 82000 | 115600 | 34793 | 49049 | 13600 | 27332 | 5770 | 11597 | 2720 | 5466 | 1154 | 2319 | 606 |
| G0053 | 14D | 0.2942 | 80800 | 23771 | 720 | 640 | 3600 | 3200 | 212 | 188 | 1059 | 941 | 10400 | 32400 | 7600 | 3060 | 9532 | 2236 | 11768 | 59400 | 102200 | 17475 | 30067 | 18400 | 16200 | 5413 | 4766 | 3680 | 3240 | 1083 | 953 | 1422 |
| G0102 | 14D | 0.4091 | 43200 | 17673 | 520 | 1800 | 2600 | 9000 | 213 | 736 | 1064 | 3682 | 7000 | 26000 | 6400 | 2864 | 10637 | 2618 | 13255 | | 71600 | | 29292 | | 19600 | | 8018 | | 3920 | | 1604 | |
| G0174 | 14D | 0.4743 | | | | | | | | | | | | | | | | | | 50668 | | 24032 | | 16932 | | 8031 | | 3386 | | 1606 | | 438 |
| G0178 | 14D | 0.2813 | 69200 | 19466 | 960 | 1840 | 4800 | 9200 | 270 | 518 | 1350 | 2588 | | | | | | | | 62400 | 63200 | 17553 | 17778 | 23400 | 31200 | 6582 | 8777 | 4680 | 6240 | 1316 | 1755 | 318 |
| G0201 | 14D | 0.3810 | 66200 | 25222 | 1800 | 1840 | 9000 | 9200 | 686 | 701 | 3429 | 3505 | 7600 | 16400 | 4000 | 2896 | 6248 | 1524 | 7772 | 95200 | 74000 | 36271 | 28194 | 28800 | 24800 | 10973 | 9449 | 5760 | 4960 | 2195 | 1890 | 294 |
| 3429 | 14D | 0.4400 | 72000 | 31680 | 720 | 720 | 3600 | 3600 | 317 | 317 | 1888 | 1888 | | | | | | | | | | | | | | | | | | | | |
| 3492 | 28D | 0.5212 | 69200 | 36067 | 240 | 2240 | 1200 | 11200 | 125 | 1167 | 625 | 5837 | 9200 | 41600 | 4800 | 4795 | 21682 | 2502 | 24184 | 90400 | 100800 | 47116 | 52537 | 29600 | 6400 | 15428 | 3336 | 5920 | 1280 | 3086 | 667 | |
| 3584 | 28D | 0.3398 | 84000 | 28543 | 1560 | 1466 | 7800 | 7332 | 698 | 498 | 2650 | 2491 | | | | | | | | 63000 | 78000 | 21407 | 26504 | 14800 | 14800 | 5029 | 5029 | 2960 | 2960 | | 1005.8 | |
| 3593 | 28D | 0.4034 | 83200 | 33563 | 1200 | 2320 | 6000 | 11600 | 484 | 936 | 2420 | 4679 | | | | | | | | | 108000 | | 43567 | | 38400 | | 15491 | | 7680 | | 3098 | |
| 3654 | 28D | 0.4753 | 64800 | 30799 | 800 | 1400 | 4000 | 7000 | 380 | 665 | 1901 | 3327 | 4932 | 51468 | 3200 | 2344 | 24463 | 1521 | 25984 | | 53400 | | 25381 | | 15000 | | 7130 | | 3000 | | 1426 | 1524 |
| 3706 | 28D | 0.3992 | 85600 | 34171 | 800 | 2106 | 4000 | 10532 | 319 | 841 | 1597 | 4204 | 4800 | 52000 | 4132 | 1916 | 20758 | 1649 | 22408 | 60668 | 61800 | 24219 | 24671 | 25468 | 11200 | 10167 | 4471 | 5094 | 2240 | 2033 | 894 | 1344 |
| 3728 | 28D | 0.2375 | 70800 | 16815 | 586 | 2186 | 2932 | 10932 | 139 | 519 | 696 | 2596 | 5400 | 47800 | 4000 | 1283 | 11353 | 950 | 12303 | 58268 | 62268 | 13839 | 14789 | 9468 | 25600 | 2249 | 6080 | 1894 | 5120 | 450 | 1216 | 533 |
| 3808 | 28D | 0.3938 | 83600 | 32922 | 960 | 1840 | 4800 | 9200 | 378 | 725 | 1890 | 3623 | 4600 | 50600 | 2400 | 1811 | 19926 | 945 | 20871 | 54000 | | 21265 | | 8600 | | 3387 | | 1720 | | 677 | | 1374 |
| G0060 | 28D | | 78400 | | 1040 | 1120 | 5200 | 5600 | | | | | 4000 | 28800 | 3600 | | | | | 72800 | 58000 | | | 15200 | 20800 | | | 3040 | 4160 | | | 957 |
| G0075 | 28D | | 82400 | | 480 | 1360 | 2400 | 6800 | | | | | 5200 | 56400 | 2600 | | | | | 61200 | 66400 | | | 35200 | 26000 | | | 7040 | 5200 | | | 426 |
| G0115 | 28D | 0.3230 | 78800 | 25452 | 1120 | 960 | 5600 | 4800 | 362 | 310 | 1809 | 1550 | 13600 | 14800 | 2000 | 4393 | 4780 | 646 | 5426 | 110600 | 79600 | 35724 | 25711 | 47400 | 13000 | 15310 | 4199 | 9480 | 2600 | 3062 | 840 | 1191 |
| G0170 | 28D | 0.4452 | 71200 | 31698 | 880 | 2640 | 4400 | 13200 | 392 | 1175 | 1959 | 5877 | 3468 | 51468 | 3200 | 1544 | 22914 | 1425 | 24338 | 77468 | 56800 | 34489 | 25287 | 42932 | 26200 | 19113 | 11664 | 8586 | 5240 | 3823 | 2333 | 992 |
| 3426 | 2D | 0.3474 | 78668 | 27329 | 3200 | 3734 | 6400 | 7468 | 1112 | 1297 | 2223 | 2594 | 2800 | 44400 | 3600 | 973 | 15425 | 1251 | 16675 | 102400 | 99868 | 35574 | 34694 | 28400 | 13600 | 9866 | 4725 | 14200 | 6800 | 4933 | 2362 | 1026 |
| 3590 | 2D | 0.4498 | | | | | | | | | | | | | | | | | | 51332 | 62000 | 23089 | 27888 | 10132 | 24800 | 4557 | 11155 | 5066 | 12400 | 2279 | 5578 | |
| 3691 | 2D | 0.6056 | 60600 | 36699 | 2600 | 3000 | 5200 | 6000 | 1575 | 1817 | 3149 | 3634 | 5468 | 67068 | 2932 | 3311 | 40616 | 1776 | 42392 | 66932 | 67468 | 40534 | 40859 | 18800 | 22668 | 11385 | 13728 | 9400 | 11334 | 5693 | 6864 | 1090 |
| 3744 | 2D | 0.3893 | 76800 | 29898 | 2400 | 5400 | 4800 | 10800 | 934 | 2102 | 1869 | 4204 | 3068 | 46932 | 4400 | 1194 | 18271 | 1713 | 19984 | 56532 | 42000 | 22008 | 16351 | 27868 | 22000 | 10849 | 8565 | 13934 | 11000 | 5425 | 4282 | |
| 3402 | 2D | 0.4097 | | | | | | | | | | | 4800 | 66400 | 4800 | 1967 | 27204 | 1967 | 29171 | 98400 | 62000 | 40314 | 25401 | 20400 | 26600 | 8358 | 10898 | 10200 | 13300 | 4179 | 5449 | 813 |
| G0006 | 2D | 0.5990 | 66800 | 40013 | 2400 | 6400 | 4800 | 12800 | 2875 | 3834 | 2875 | 7667 | | | | | | | | 92532 | | 55427 | | 19068 | | 11422 | | 9534 | | 5711 | | 1896 |
| G0118 | 2D | 0.4434 | 79200 | 35114 | 2400 | 5400 | 4800 | 10800 | 1064 | 2394 | 2128 | 4788 | 4400 | 50200 | 3200 | 1951 | 22257 | 1419 | 23675 | | 66200 | | 29350 | | 31400 | | 13922 | | 15700 | | 6961 | 1646 |
| G0133 | 2D | | 72400 | | 2000 | 2734 | 4000 | 5468 | | | | | 13320 | 36532 | 5200 | | | | | 43400 | 62800 | | | 8400 | 35000 | | | 4200 | 17500 | | | 1563 |
| G0136 | 2D | 0.4912 | 70800 | 34777 | 1466 | 1466 | 2932 | 2932 | 720 | 720 | 1440 | 1440 | 7200 | 61200 | 6000 | 3537 | 30061 | 2947 | 33009 | 66400 | 72400 | 32616 | 35563 | 25600 | 26800 | 12575 | 13164 | 12800 | 13400 | 6287 | 6582 | 1479 |
| G0192 | 2D | 0.4510 | 72800 | 32833 | 2800 | 3600 | 5600 | 7200 | 1263 | 1624 | 2526 | 3247 | 11600 | 47468 | 4800 | 5232 | 21408 | 2165 | 23573 | | | | | | | | | | | | | 516 |
| 3683 | 4D | 0.5654 | 64400 | 36412 | | | | | | | | | 10800 | 33600 | 4400 | 6106 | 18997 | 2488 | 21485 | 101400 | 67332 | 57332 | 38069 | 19200 | 7068 | | | 4800 | 1767 | 2714 | 999 | 561 |
| 3722 | 4D | 0.4963 | 74800 | 37123 | 1050 | 2150 | 4200 | 8600 | 521 | 1067 | 2084 | 4268 | 4000 | 58000 | 6400 | 1985 | 28785 | 3176 | 31962 | 62400 | 46268 | 30969 | 22963 | 25800 | 15200 | 12805 | 7544 | 6450 | 3800 | 3201 | 1886 | 678 |
| G0022 | 4D | 0.4567 | 68800 | 31421 | 1033 | 1800 | 4132 | 7200 | 472 | 822 | 1887 | 3288 | 8400 | 24400 | 2000 | 3836 | 11143 | 913 | 12057 | 41332 | 48800 | 18876 | 22287 | 13200 | 7600 | 6028 | 3471 | 3300 | 1900 | 1507 | 868 | 1371 |
| G0026 | 4D | 0.5243 | 87200 | 45719 | 1650 | 2125 | 6600 | 8500 | 865 | 1114 | 3460 | 4457 | 4800 | 55800 | 2600 | 2517 | 29256 | 1363 | 30619 | 57600 | 95868 | 30200 | 50264 | 17200 | 30532 | 9018 | 16008 | 4300 | 7633 | 2254 | 4002 | 3888 |
| G0030 | 4D | 0.3717 | 66800 | 24830 | 933 | 1100 | 3732 | 4400 | 347 | 409 | 1387 | 1635 | 5600 | 24000 | 2400 | 2082 | 8921 | 892 | 9813 | 59600 | 54000 | 22153 | 20072 | 12800 | 11000 | 4758 | 4089 | 3200 | 2750 | 1189 | 1022 | 4763 |
| G0031 | 4D | 0.4152 | 78400 | 32552 | 1300 | 800 | 5200 | 3200 | 540 | 332 | 2159 | 1329 | 6000 | 30000 | 8400 | 2491 | 12456 | 3488 | 15944 | 50132 | 90800 | 20815 | 37700 | 9068 | 21000 | 3765 | 8719 | 2267 | 5250 | 941 | 2180 | 2508 |
| G0045 | 4D | 0.4246 | 76400 | 32439 | 1350 | 2850 | 5400 | 11400 | 573 | 1210 | 2293 | 4840 | 6800 | 37600 | 2600 | 2887 | 15965 | 1104 | 17069 | 92400 | 60200 | 39233 | 25561 | 19800 | 4400 | 8407 | 1868 | 4950 | 1100 | 2102 | 467 | 1695 |
| G0094 | 4D | | 77600 | | 1567 | 2700 | 6268 | 10800 | | | | | | | | | | | | 82600 | 88600 | | | 8600 | 9800 | | | 2150 | 2450 | | | |
| G0186 | 4D | 0.4794 | 77600 | 37201 | 1533 | 2200 | 6132 | 8800 | 735 | 1055 | 2940 | 4219 | | | | | | | | 55800 | 66200 | 26751 | 31736 | 19800 | 23200 | 9492 | 11122 | 4950 | 5800 | 2373 | 2781 | 450 |
| G0191 | 4D | 0.3509 | 70000 | 24563 | 700 | 2100 | 2800 | 8400 | 246 | 737 | 983 | 2948 | 5600 | 23200 | 3600 | 1965 | 8141 | 1263 | 9404 | 50400 | 71400 | 17685 | 25054 | 15600 | 23400 | 5474 | 8211 | 3900 | 5850 | 1369 | 2053 | |
| 3599 | 7D | 0.4879 | 74000 | 36105 | 960 | 1520 | 4800 | 7600 | 468 | 742 | 2342 | 3708 | 4800 | 36800 | 2400 | 2342 | 17955 | 1171 | 19126 | 56400 | 79800 | 27518 | 38934 | 8000 | 16800 | 3903 | 8197 | 1600 | 3360 | 781 | 1639 | 203 |
| 3632 | 7D | | 66400 | | 960 | 1680 | 4800 | 8400 | | | | | 7600 | 55400 | 2000 | | | | | 70000 | 55800 | | | 28800 | 10800 | | | 5760 | 2160 | | | 692 |
| 3665 | 7D | 0.3835 | 60400 | 23163 | 480 | 880 | 2400 | 4400 | 184 | 337 | 920 | 1687 | 6000 | 28800 | 7200 | 2301 | 11045 | 2761 | 13806 | | 71600 | | 27459 | | 16000 | | 6136 | | 3200 | | 1227 | 399 |
| 3746 | 7D | 0.5075 | 66400 | 33698 | 560 | 1720 | 2800 | 8600 | 284 | 873 | 1421 | 4365 | 5200 | 42400 | 4400 | 2639 | 21518 | 2233 | 23751 | 84400 | 69400 | 42833 | 35220 | 22200 | 8000 | 11267 | 4060 | 4440 | 1600 | 2253 | 812 | 587 |
| G0016 | 7D | 0.4869 | 86400 | 42068 | 880 | 1280 | 4400 | 6400 | 428 | 623 | 2142 | 3116 | | | | | | | | 31200 | 72400 | 15191 | 35252 | 10800 | 39200 | 5259 | 19086 | 2160 | 7840 | 1052 | 3817 | |
| G0055 | 7D | 0.3485 | 71200 | 24813 | 640 | 840 | 3200 | 4200 | 223 | 293 | 1115 | 1464 | 5200 | 25600 | 3600 | 1812 | 8922 | 1255 | 10176 | 113000 | 108800 | 39381 | 37917 | 25400 | 26000 | 8852 | 9061 | 5080 | 5200 | 1770 | 1812 | 1032 |
| G0069 | 7D | 0.3493 | 74400 | 25988 | 586 | 1360 | 2932 | 6800 | 205 | 475 | 1024 | 2375 | 4120 | 33160 | 7600 | 1439 | 11583 | 2655 | 14237 | 58400 | 82400 | 20399 | 28782 | 20400 | 28400 | 7126 | 9920 | 4080 | 5680 | 1425 | 1984 | 863 |
| G0172 | 7D | 0.2869 | | | | | | | | | | | 6000 | 20600 | 3600 | 1721 | 5910 | 1033 | 6943 | 50800 | 46200 | 14575 | 13255 | 13200 | 20400 | 3787 | 5853 | 2640 | 4080 | 757 | 1171 | 417 |
| G0179 | 7D | 0.3004 | 70400 | 21148 | 1680 | 1560 | 8400 | 7800 | 505 | 469 | 2523 | 2343 | 7600 | 22200 | 3600 | 2283 | 6669 | 1081 | 7750 | 49200 | | 14780 | | 13200 | | 3965 | | 2640 | | 793 | | 345 |
